# Differences in orthographic processing across species identified by a transparent computational model

**DOI:** 10.1101/2024.06.25.600635

**Authors:** Benjamin Gagl, Ivonne Weyers, Susanne Eisenhauer, Christian J. Fiebach, Janos Pauli, Michael Colombo, Damian Scarf, Johannes C. Ziegler, Jonathan Grainger, Onur Güntürkün, Jutta L. Mueller

**Affiliations:** Self-learning Systems Lab, University of Cologne, Cologne, Germany; Department of Linguistics, University of Vienna, Vienna, Austria; Basque Center on Cognition, Brain and Language, Donostia-San Sebastián, Spain; Department of Psychology, Goethe University Frankfurt, Frankfurt am Main, Germany; Department of Psychology, University of Otago, Dunedin, New Zealand; Aix-Marseille University, CNRS, Aix-en-Provence, France; Department of Biopsychology, Ruhr University Bochum, Bochum, Germany; Vienna Cognitive Science Research Hub, Vienna, Austria

## Abstract

The ability to robustly recognize strings of letters, a cornerstone of reading, was observed in baboons and pigeons despite their lack of phonological and semantic knowledge. Here, we apply a comparative modeling approach to investigate the neuro-cognitive basis of orthographic decision behavior in humans, baboons, and pigeons, addressing whether phylogenetic relatedness entails similar underlying neuro-cognitive phenotypes. We use the highly transparent SpeechLess Reader (SLR) model, which assumes letter-string recognition based on a computational implementation of predictive coding, whereby orthographic decisions rely on prediction-error signals emerging from multiple representational levels: visual-pixel, letter, and letter-sequence representations. We investigate which representations species use during successful orthographic decision-making. We introduce multiple SLR variants, each including one or more prediction-error representations, and compare the simulations of each variant with orthographic decisions from individuals of three species after learning letter strings without meaning. Humans predominantly relied on letter-sequence-level representations, resulting in the highest task performance in behavior and model simulations. Baboons also relied on sequence-based representations, but in combination with pixel- and letter-level representations. In contrast, pigeons relied more on pixel and letter-level representations. These findings suggest that the orthographic representations used in orthographic decisions reflect phylogenetic distance: Humans and baboons use more similar representations than pigeons. Overall, the description of orthographic decisions based on a small set of representations and computations was highly successful in describing behavior, even for humans who mastered reading in its entirety.

## Introduction

Script is a hallmark of human cultural evolution unmatched in the animal kingdom [1]. Reading is considered exclusive to humans, requiring explicit instruction and specialized cognitive skills to retrieve word representations and meaning [2–5]. Biological foundations of reading ability have been described, for instance, in the finding of a visual word form area in the human left-ventral occipito-temporal cortex [1, 6] or of genetic risk for dyslexia [7]. The description of the biological basis of reading and reading-like behavior for baboons [8] and pigeons [9] raises the question of the evolutionary roots of reading ability. Both species learned to distinguish remembered orthographic stimuli (i.e., similar to written words) from novel letter combinations in orthographic decision tasks. While this suggests similar skills in non-human animals and humans at the surface, how the task is solved across species, i.e., which cognitive processes were involved, remains an open question. Note that computer vision models with drastically different architectures can also solve similar tasks [10–13], indicating that there are many ways to solve orthographic processing tasks. In the current study, we investigate the cognitive basis of reading(-like) behavior (i.e., orthographic decisions) in humans, baboons, and pigeons and test if phylogenetic proximity leads to more human-like cognitive processes in primates than in birds.

Humans typically learn to read through grapheme-phoneme conversion only once robust phonological and semantic representations are already in place [14–16], thereby aligning their reading rate to that of speech in fluent readers [17]. Still, on the level of word recognition, orthographic word characteristics have been identified as central aspects for the differentiation between word and non-word letter strings (i.e., lexical decisions; Ref. [18–22]). Similar orthographic decision tasks, without the involvement of explicit semantic processing, have been implemented in baboons [8], pigeons [9], humans [23] and Computer vision models [10, 11] (see also, e.g., [24–26] for learning paradigms also implementing phonology and semantic information in humans).

Here, we investigate the representations underlying orthographic decisions in humans, baboons, and pigeons. To implement this, we use *neuro-cognitive computational phenotyping*, an approach that leverages transparent computational models, previously successful in psychiatry and personality psychology [27–29]. At its core, this approach (i) describes which neuro-cognitive operations are involved in decisions and (ii) allows a comparison of the types of representation implemented across individuals, and therefore also across species. Hence, we investigate the evolutionary roots of reading-like behavior using computational neurocognitive phenotypes by comparing species that differ in phylogenetic proximity.

### The Speechless Reader Model (SLR)

We developed the SLR combining neuro-cognitive assumptions of visual and orthographic processing in human visual word recognition [6, 20, 21, 30] constrained by the principles of predictive coding [31–33]. Predictive coding is a theory of neuronal functioning that describes how the brain implements efficient information processing while respecting the energy-consumption constraints of the brain [34]. The core of the assumption is the reduction of sensory bottom-up information flow by predicting away the expected information of the stimulus (e.g., see examples from vision [21, 31, 35], or language processing [36–38]). The current SLR implementation models orthographic decisions based on visual and orthographic representations, optimizing processing based on knowledge predictions (i.e., based on lexicon entries only without explicit, classically investigated context-based predictions; see [21] for a detailed discussion). The SLR neglects phonological and semantic representations, while capturing the visual-orthographic processes involved in the lexical decision tasks. Thus, the SLR is, by design, optimally suited to simulate orthographic decisions in non-human species (here: baboons and pigeons) naive to phonology and semantics, with a minimal set of assumptions. For humans, simulating orthographic decisions based solely on visual-orthographic information enables us to assess the extent to which it explains variance.

The core of the model consists of three types of prediction error representations. All integrate the model input (i.e., the visually presented letter string) with the knowledge stored in the lexicon of each participant (i.e., the stored words), which, in turn, differs across participants depending on experience or learning history. For example, the orthographic prediction error (oPE; Ref. [20, 21]) integrates word knowledge and sensory input at the pixel level to reduce neuronal processing effort when representing sensory input. The oPE is a simple demonstration of how predictive coding principles can be applied to early stages of visual word recognition, calculating a prediction error by subtracting expected visual information (i.e., implemented by a pixel-by-pixel mean of the 15% most frequent words in the lexicon; Ref. [20]) from the actual sensory input encountered while processing a written word. This purely visual, pixel-level oPE is a well-established orthographic word characteristic (i.e., [20, 21]). Thus, this simple pixel-level operation produces an abstract and efficient representation of the word or letter string, supporting orthographic decisions.

Here, to extend the model’s capabilities to higher-level (i.e., more abstract) representations, we applied the same principles (i.e., derived from predictive coding) to letters and letter sequences. We calculated the probabilities of letter occurrence at any given position and of letter sequences, and used these to generate predictions and, in the next step, prediction errors specific to the model’s respective processing level. For example, in English, the letter ”s” is highly likely to appear at the end of a word, marking the plural (see [37] for similar ideas in models implemented for speech recognition). For the letter sequence representation, we, again, apply the same logic, starting with the first letter, such that we generate predictions based on the probability of the sequence given the letter strings in the lexicon (e.g., how probable is ”Ha” or ”Hau” given the lexicon). Thus, high-level letter-sequence prediction errors represent unexpected sequences given the stored items in the lexicon.

The current version of the SLR thus combines information from three prediction error representations that operate at three distinct levels (see Fig. 1): pixel (oPE), letter (LPE), and letter-sequence (sPE). In the full model (i.e., the computational phenotype including all three types of representations), prediction error values are calculated for all three levels, which are summed up to generate a word-likeness estimate for each encountered letter string. This estimate then forms the basis for the orthographic decision (see Eq. 1, 2, 3, 10, 5, and 6).

**Fig 1.**
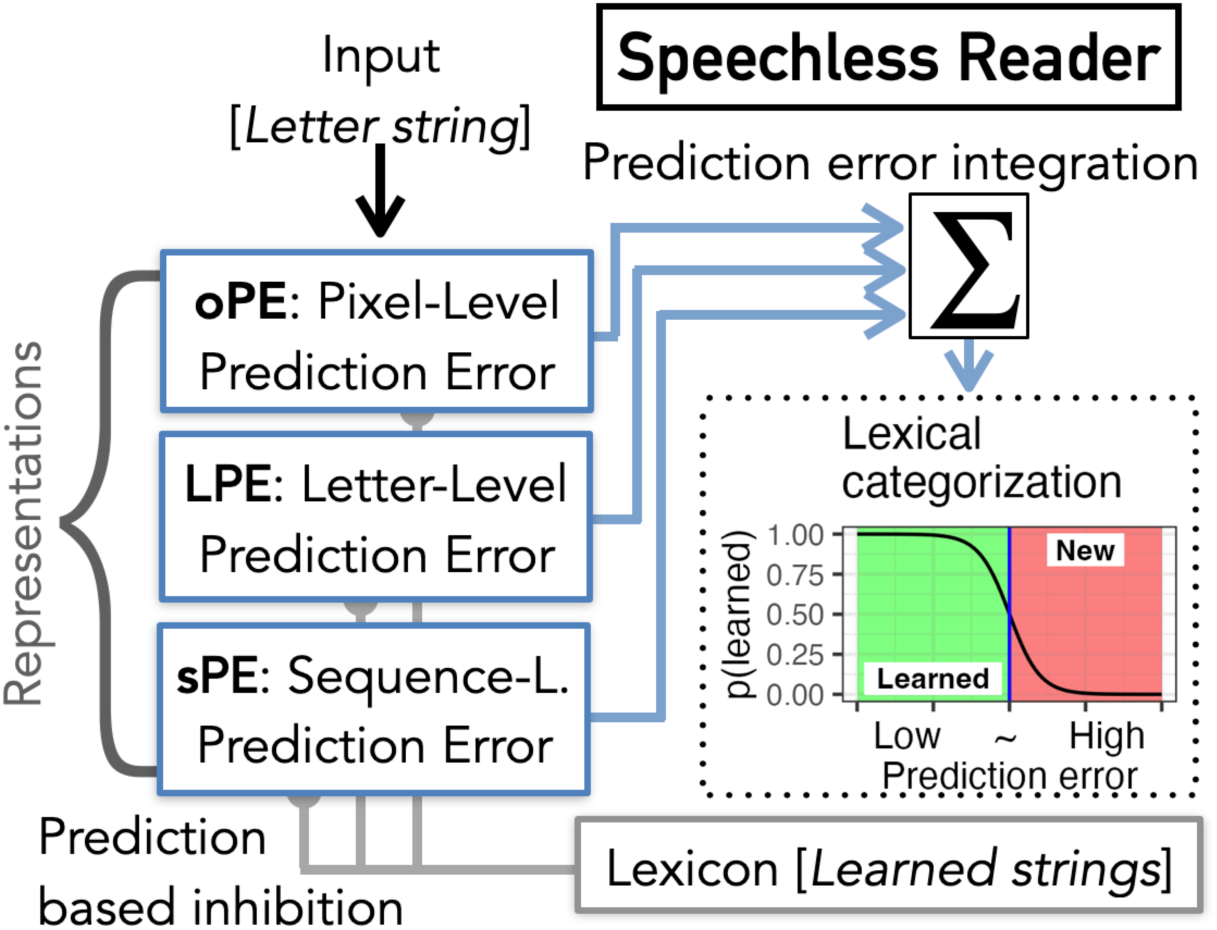
*Speechless Reader Model*, including model input and three types of prediction error representations based on: Pixel-Level visual-orthographic representations (oPE; Ref. [20, 21]), Letter-Level letter-position representations (LPE), and Sequence-Level letter-sequence representations (sPE; Ref. [30]). Prediction errors are calculated based on the learned letter strings stored in the lexicon. The resulting representations are summed into a word-likeness estimate [6]. The output of the categorization process is binary, indicating whether the input is considered learned or new (e.g., like word vs. non-word in a classical lexical decision task).

To investigate the evolutionary roots of reading-like behavior, we utilize the SLR to model three datasets: (i) human (N = 37; [23, 39]), (ii) baboon (N = 6; [8]), and (iii) pigeon data (N = 4; [9]), all learn to differentiate between learned and novel letter strings without explicit semantic and phonological associations. We created an individualized lexicon comprising the letter strings successfully learned by each human, baboon, and pigeon. We then generated model simulations of seven different model variants (i.e., similar to a digital twin; e.g., see [40]) representing all possible combinations of the three representational levels (prediction error types; e.g., using only the letter-level LPE or combining oPE and LPE). In addition, we used fitted categorization thresholds for each individual (cf. blue line in Fig. 1; see also Eq. 6; the only free parameter of the model). Then, we compared the model and participant performance for each participant (see Appendix B, Fig. S2-4), and the simplest model with the lowest mean squared error was considered the best-fitting model. For interpretation, we focus on the representations implemented in the models with the highest behavioral similarity to each participant. We consider this set of representations the neuro-cognitive phenotype for orthographic decisions of that individual.

## Results

### Model representations

First, we investigate model simulations, focusing on representations of prediction error. These representations result from integrating the sensory input of each learned and novel letter string and the prediction based on the individual lexicon from each reader (see Eq. 1 for the prediction estimation and Eq. 2 for the prediction error estimation; see Table 1 for number of items in the lexicon). As a result, the variability in the prediction error across participants reflects individual learning histories. As expected, we find reduced prediction errors on all three representational levels and all three species for learned compared to novel letter strings (see Table 2 and Fig. 2a-c,h-j,o-q), indicating that the prediction errors are indeed specifically reduced for learned words and, therefore, can be utilized to solve a learned/novel categorization task. Note that, although the differences can be relatively small, the high number of trials results in a highly exact estimation of means such that even a small difference becomes statistically significant. Thus, considering the effect size is advised.

**Fig 2.**
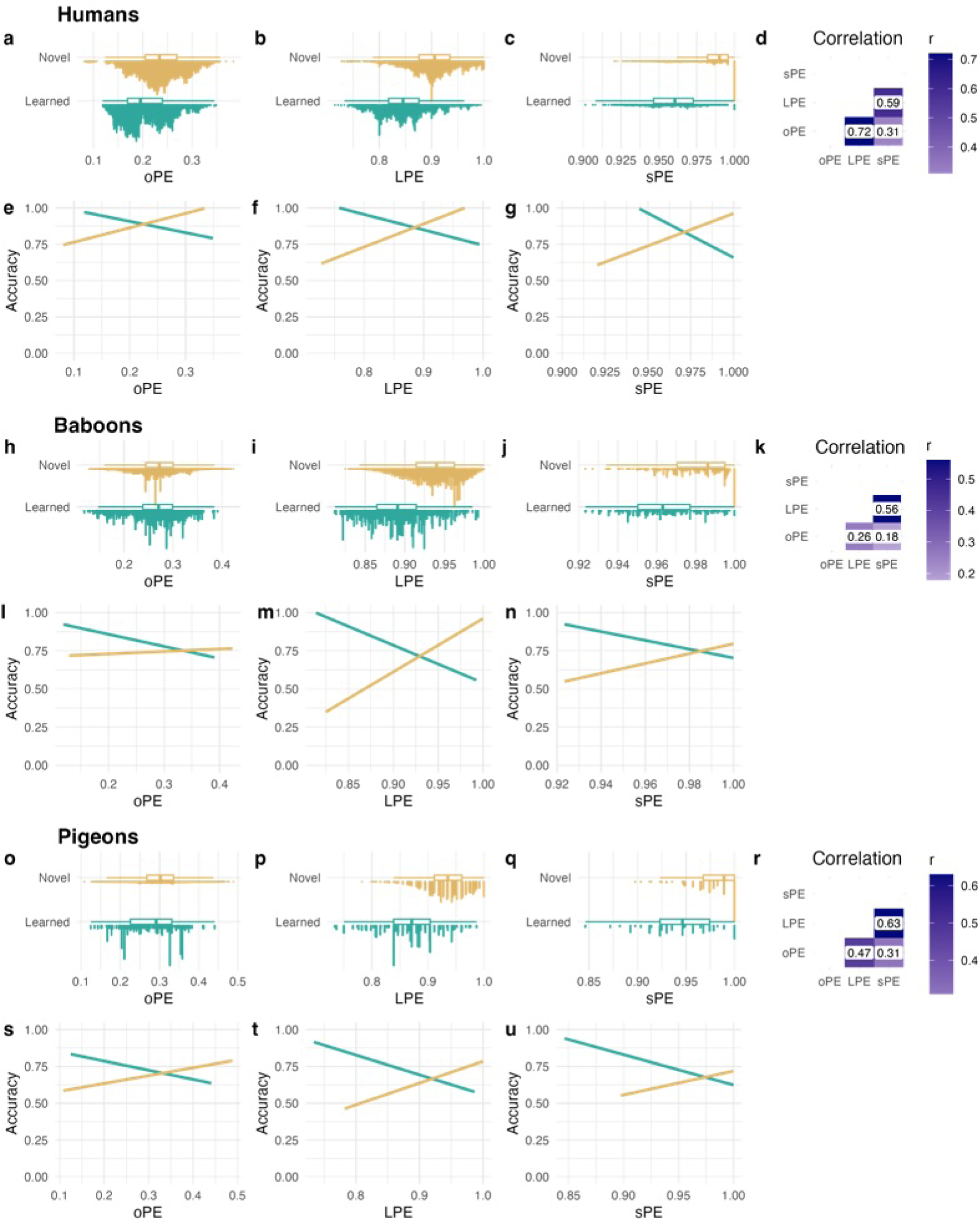
Estimated normalized prediction error representations on the pixel, letter, and letter-sequence level and their relation to human, baboon, and pigeon orthographic decision performance. (a,h,o) oPE, (b,i,p) LPE, and (c,j,q) sPE are estimated based on the individual lexica from each participant and normalized. Here, we divided them into learned (turquoise) and novel (dark yellow) letter strings (y-axis). (d,k,r) Correlation matrix comparing the three prediction error representations. (e,l,o) oPE, (f,m,t) LPE, and (g,n,u) sPE effects on decision accuracy, including lines for learned (turquoise) and novel (dark yellow) letter strings.

**Table 1.** Number of individuals and descriptive statistics of the number of items in individual lexicons for each species.

|  | N | Median | Min | Max |
| --- | --- | --- | --- | --- |
| Humans | 37 | 54 | 34 | 59 |
| Baboons | 6 | 86.5 | 67 | 271 |
| Pigeons | 4 | 24.5 | 8 | 32 |

**Table 2.**
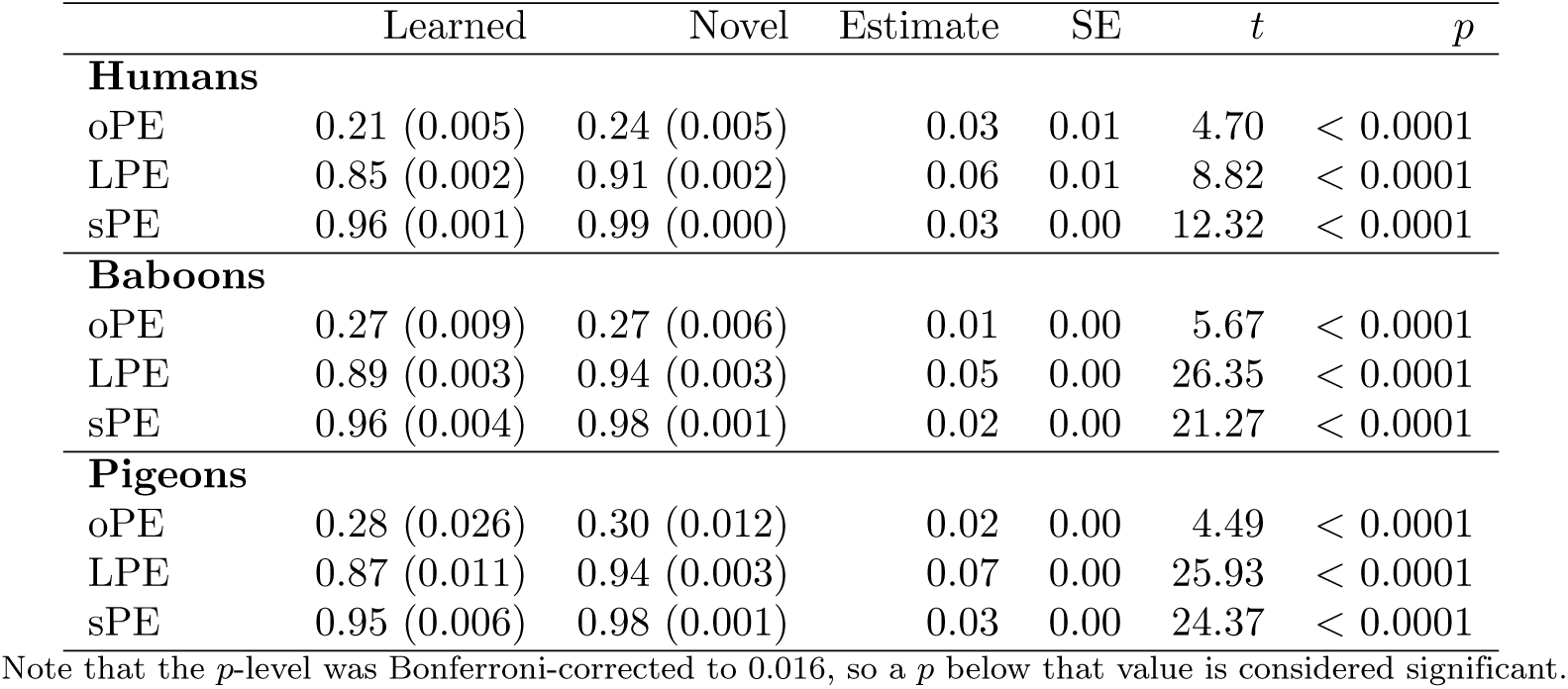
Linear mixed regression models investigating the prediction error differences between learned and novel stimuli for pixel, letter, and letter-sequence representations based on individual lexicons.

When correlating the model representations within each species, the first differences emerge. In humans, LPE and oPE show the highest correlation (Fig. 2d). For baboons and pigeons, we found the highest correlation between LPE and sPE (Fig. 2d,r). This indicates that the lower-level representations were similar across humans, whereas in baboons and pigeons letter- and letter-sequence-level representations were more similar.

### Human behavior

Mean orthographic decision accuracy showed that humans performed best, with an average accuracy higher than 90% (see Fig. 3a,b). As described before in the context of real words [20, 21], for known letter strings, we observe fewer errors and faster reaction times when prediction errors are low. We found the inverted pattern for novel letter strings, for pixel, letter, and letter-sequence representations, indicated by the significant interaction of the prediction errors and lexicality (Accuracy: see Fig. 2e-g and Table 3; for response times see Appendix A, Table S1, Fig. S1). Overall, we found the smallest difference in the oPE and the largest in the sPE.

**Fig 3.**
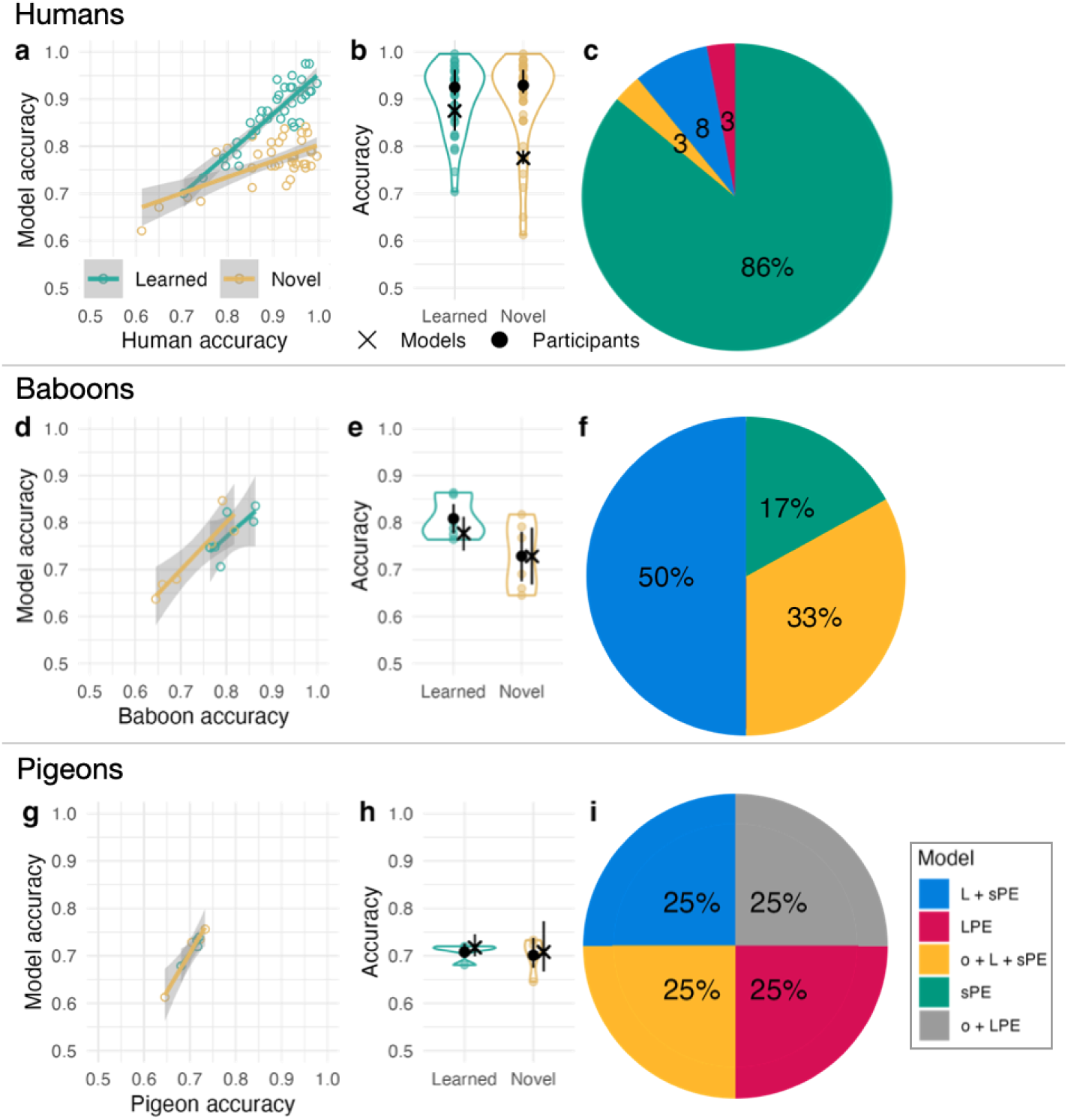
Model fit and neuro-cognitive phenotypes for decision behavior for each participant separated by species. (a,d,g) Correlation of behavioral accuracy (x) and model accuracy (y) of each participant for learned (green) and novel (yellow) stimuli. Mean model accuracy was extracted from the models that best described performance at the individual level. Lines represent linear regression lines with confidence intervals. (b,e,h) Behavioral results (mean accuracy per individual; dots and violin, separated for learned and novel words), including group mean (black dot) and the mean model simulation results of the individual winning models (black X) with bootstrapped 95% confidence intervals. Note that only the data on learned letter strings were available for pigeons. (c,f,i) Proportion of model variants that best described human (33 showing the sPE, tow the L + sPE, one the o + L + sPE, and one the LPE phenotype), baboon (three showing the L + sPE, tow the o + L + sPE, and one the sPE phenotype), and pigeon orthographic decision behavior (one showing the L + sPE, one the o + L + sPE, one the LPE, and one the o + LPE phenotype; see Appendix C, Table S2, for a full list for each participant).

**Table 3.** Generalized linear mixed regression models investigating the interaction of three prediction error representations and letter string type (Learned/Novel) in participant behavior (Response accuracy).

|  | Estimate | Std. Error | *z* | *p* |
| --- | --- | --- | --- | --- |
| **Humans** |  |  |  |  |
| oPE | 52.1 | 4.6 | 11.2 | < 0.0001 |
| Learned/Novel | 13.6 | 1.1 | 12.1 | < 0.0001 |
| oPE X Learned/Novel | -64.4 | 5.2 | -12.4 | < 0.0001 |
| LPE | 107.7 | 4.7 | 23.0 | < 0.0001 |
| Learned/Novel | 111.6 | 5.2 | 21.3 | < 0.0001 |
| LPE X Learned/Novel | -129.6 | 6.0 | -21.6 | < 0.0001 |
| sPE | 221.3 | 5.4 | 41.1 | < 0.0001 |
| Learned/Novel | 245.2 | 7.0 | 35.3 | < 0.0001 |
| sPE X Learned/Novel | -253.4 | 7.1 | -35.7 | < 0.0001 |
| **Baboons** |  |  |  |  |
| oPE | 1.6 | 0.3 | 5.1 | < 0.0001 |
| Learned/Novel | 3.4 | 0.2 | 18.0 | < 0.0001 |
| oPE X Learned/Novel | -10.9 | 0.7 | -15.6 | < 0.0001 |
| LPE | 20.0 | 0.3 | 63.2 | < 0.0001 |
| Learned/Novel | 37.3 | 0.5 | 78.7 | < 0.0001 |
| LPE X Learned/Novel | -40.3 | 0.5 | -76.1 | < 0.0001 |
| sPE | 19.5 | 0.6 | 31.6 | < 0.0001 |
| Learned/Novel | 50.1 | 0.5 | 98.0 | < 0.0001 |
| sPE X Learned/Novel | -50.9 | 0.5 | -96.8 | < 0.0001 |
| **Pigeons** |  |  |  |  |
| oPE | 2.6 | 0.3 | 7.6 | < 0.0001 |
| Learned/Novel | 2.4 | 0.2 | 10.5 | < 0.0001 |
| oPE X Learned/Novel | -7.8 | 0.8 | -10.0 | < 0.0001 |
| LPE | 6.8 | 0.4 | 17.1 | < 0.0001 |
| Learned/Novel | 13.9 | 0.6 | 23.2 | < 0.0001 |
| LPE X Learned/Novel | -15.4 | 0.7 | -22.8 | < 0.0001 |
| sPE | 7.5 | 0.8 | 9.3 | < 0.0001 |
| Learned/Novel | 15.6 | 1.4 | 11.4 | < 0.0001 |
| sPE X Learned/Novel | -16.0 | 1.4 | -11.2 | < 0.0001 |

Note that the *p*-level was Bonferroni-corrected to 0.016, so a *p* below that value is considered significant.

### Baboon and pigeon behavior

Baboons and pigeons performed above chance level, with mean accuracies of approximately 75% for baboons (see 3d,e) and just above 70% for pigeons (see 3g,h). As in humans, orthographic decision accuracy in both species showed a significant interaction between prediction error and stimulus type (learned vs. novel) across all three representational levels (oPE, LPE, and sPE; see Table 3 and Fig. 2l-n,s-u).

### Model behavior

The model simulations mirror this general pattern of accuracy results (see Fig. 3a,b,d,e,g,h). Thus, on average, the best-fitting speechless reader models performed similarly to baboons and pigeons but were less accurate than humans, even though the model simulations for humans resulted in the highest accuracies.

Nevertheless, our model simulations were highly correlated with behavior across all three species (see Table 4 and Figure 3a,d,g), indicating that the selected models adequately represent the behavior of the individual participants. The pie charts in Figure 3 show which model variants (including different prediction-error representations) achieved the highest fit for the individual human, baboon, and pigeon data, respectively. We found that, for humans, the best-fitting model included letter-sequence-level representations in all but one participant. For 86% of participants, the best-fitting model used exclusively letter-sequence-level representations (sPE). For 8%, it consisted of a combination of letter-sequence-level and letter-level representations (L + sPE), and for 3%, the whole model with all three representations was most adequate (o + L + sPE). The behavior of one participant (3%) without the letter-sequence-level representation was best explained by a model with only the letter-level representation (LPE).

**Table 4.** Linear regression models investigating the association of model simulations and participant behavior.

| Species ($R^2$) | Parameter | Est. (Std. Error) | $t$ | $p$ |
| --- | --- | --- | --- | --- |
| Humans (0.58) | Model | 1.09 (0.11) | 9.8 | < 0.001 |
|  | Learned/Novel | 0.10 (0.02) | 6.1 | < 0.001 |
| Baboons (0.85) | Model | 0.78 (0.15) | 5.3 | < 0.001 |
|  | Learned/Novel | -0.04 (0.02) | -2.2 | 0.056 |
| Pigeons (0.95) | Model | 0.61 (0.05) | 11.5 | < 0.001 |
|  | Learned/Novel | 0.01 (0.01) | 0.3 | 0.799 |

Note. Learned/Novel refers to the main effect of stimulus type. We tested the interaction between stimulus conditions and model simulation and found no significant interaction in either humans or baboons.

For baboons, the same model variants (i.e., the same set of representation combinations) exhibited the best fit, albeit in different portions. Only the behavior of one baboon was best described by the model variant, which was most common in humans, using only the sPE representation. Still, all baboons included the sequence level representation in the phenotype. Most baboons (50%; N = 3) required the implementation of a combination of letter-sequence-level and letter-level representations (L + sPE), the second most common type of representation found in humans.

For pigeons, the set of the best-fitting models was different. All models included letter-level representations (LPE). One pigeon used all three representations (o + L + sPE), one pixel and letter-level representations (o + LPE), one letter and letter-sequence-level representations (L + sPE), and one pigeon only used the letter-level representations (LPE). Please find an individual-level error analysis in Appendix C, Table S2.

To measure the stability of the phenotyping, we estimated the split-half reliability (Table 5 upper section). We based the analysis on the agreement between the representations from the individual models fitted to one-half of the data and those fitted to the other half. We implemented random splitting and repeated the procedure one hundred times. Overall, reliability was high (agreement *>* 65% for all representations), with the results for the letter and letter-sequence representations in humans showing the lowest reliability (69.9%). To assess whether the phenotypic estimates rely heavily on the best-fitting model, we reevaluated the three representations using the mean squared error across all implemented models (see the lower section of Table 5). The lowest mean-squared error emerged when models for humans included letter-sequence-level representations. For baboons, we find letter-sequence-level and letter-level representations showing the lowest errors. For pigeons, the error is lowest for models that include the letter-level prediction error. These analyses substantiate the main findings from the best-fitting models.

**Table 5.**
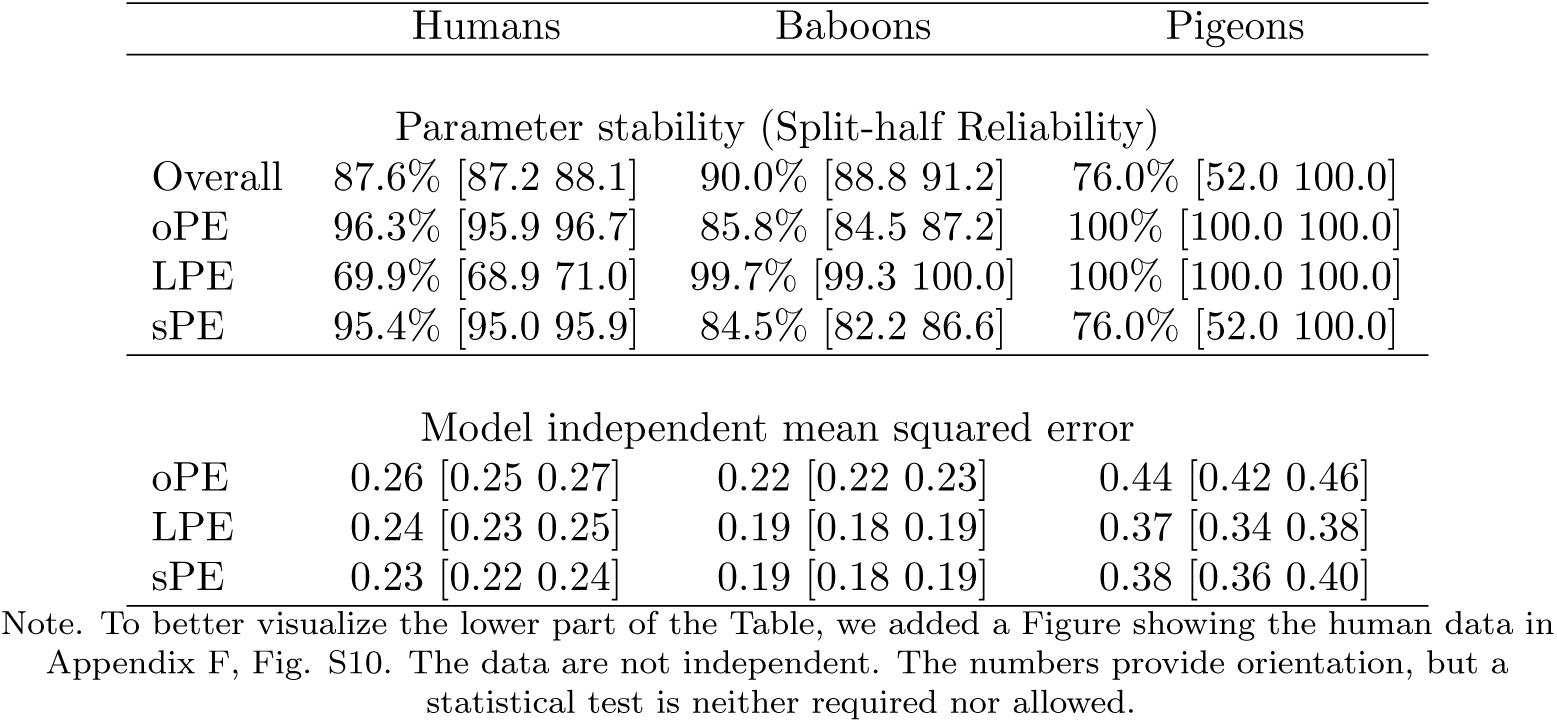
Investigation of the reliability and stability of the individual phenotype models. Upper section: Mean split-half reliability based on representation agreement in percent, including 95% confidence intervals in parentheses. Splitting was implemented randomly and repeated 100 times. Lower section: Mean squared errors, including 95% confidence intervals, from all implemented models (i.e., for all seven representation combinations and all thresholds) separated by species and representation (i.e., combining models that included the representation; for the oPE measure, this would include o + L + sPE, o + LPE, o + sPE, and oPE models). The latter part is included to investigate the independence of the phenotype estimation from the individual best-fitting model.

|  | Humans | Baboons | Pigeons |
| --- | --- | --- | --- |
| Parameter stability (Split-half Reliability) |  |  |  |
| Overall | 87.6% [87.2 88.1] | 90.0% [88.8 91.2] | 76.0% [52.0 100.0] |
| oPE | 96.3% [95.9 96.7] | 85.8% [84.5 87.2] | 100% [100.0 100.0] |
| LPE | 69.9% [68.9 71.0] | 99.7% [99.3 100.0] | 100% [100.0 100.0] |
| sPE | 95.4% [95.0 95.9] | 84.5% [82.2 86.6] | 76.0% [52.0 100.0] |
| Model independent mean squared error |  |  |  |
| oPE | 0.26 [0.25 0.27] | 0.22 [0.22 0.23] | 0.44 [0.42 0.46] |
| LPE | 0.24 [0.23 0.25] | 0.19 [0.18 0.19] | 0.37 [0.34 0.38] |
| sPE | 0.23 [0.22 0.24] | 0.19 [0.18 0.19] | 0.38 [0.36 0.40] |

Note. To better visualize the lower part of the Table, we added a Figure showing the human data in Appendix F, Fig. S10. The data are not independent. The numbers provide orientation, but a statistical test is neither required nor allowed.

## Discussion

In an orthographic decision task (i.e., categorization of letter strings as learned or novel), we used a model-based approach to infer the representations used for orthographic decisions in humans, baboons, and pigeons, based on a transparent computational account: the Speechless Reader model. Previous investigations focused on behavioral performance similarities across species [8, 9], which are also replicated here (e.g., similar behavioral pattern of prediction error effect). In contrast, the present study focuses on investigating differences between species and individuals using neurocognitive computational phenotyping [28, 29]. We integrate results from humans, baboons, and pigeons to compare the representations used for orthographic decisions of species separated by 324 million years of independent evolution [41]. Overall, we found no evidence that any single representation is species-specific, but one combination of representations is specific to pigons and shows quantitative differences in the frequency of phenotypes within a species.

Although the lexicon size differed between species after learning, learned words elicited lower prediction error estimates at all levels of representation (for a systematic investigation of phenotypes while learning, see [42]). When correlated with behavior for all species and representations, the previously described characteristic interaction pattern was found (i.e., positive prediction error effect for learned strings and negative effect for novel letter strings, see [20, 21]). Yet, the contributions of those representational levels to the task performance varied between species. Humans predominantly relied on letter-sequence-level representations as the basis for their orthographic decisions, suggesting that they extract relevant information from letter strings based on combinations of at least two letters. To some extent, this is also true for baboons. However, they used letter-level representations to a greater extent than humans do. For only one baboon, we found the phenotype typical for humans (i.e., letter-sequence-level representation only; found in 86% of humans).

In pigeons, orthographic decisions were compatible with models incorporating multiple representational levels, with substantial inter-individual variability. Unlike humans, pigeons did not consistently require letter-sequence-level representations to account for their behavior, and no single representational profile characterized the species as a whole. Importantly, model-independent analyses showed that representations including letter-position and letter-sequence information accounted for pigeon behavior equally well, whereas pixel-level representations alone provided a poorer fit. Given the small sample size (N = 4), strong species-level claims about representational preferences are unwarranted. Instead, the results suggest that pigeons can solve orthographic decision tasks using flexible combinations of representational levels, without systematic reliance on higher-order sequence representations. More generally, the observed variability contrasts with the more homogeneous reliance on sequence-level representations in humans and supports a graded view in which species differ in the consistency with which higher-order orthographic representations are engaged.

The perceptual demands of the analyzed orthographic decision studies, which mirror the tasks of local and global visual processing, can help explain the pattern differentiating between species. Comparative studies involving humans, non-human primates, and pigeons have revealed that perceptual specializations on global or local stimulus features impact various cognitive tasks [43–46]. The striking similarity of our findings to those on global/local processing suggests a potential overlap in the visuo-cognitive processes required for visual orthographic integration. Humans have a strong propensity to group small elements into global configurations. Thus, they preferably process a large letter instead of its constituent small letters [47], easily see the global shape from partially deleted line drawings [48], integrate distant patterns into illusory shapes [49], and exploit perceptual principles like similarity and proximity to see a global form [50]. Baboons respond more slowly to global than to local targets [51], while Macaques selectively attend to fine details of compound stimuli [52]. Detailed analyses of these and similar results suggest that these differences occur at the attentional level, where perceptual grouping operations seem more attention-demanding for monkeys. Attentional differences in human early visual cortical areas may be especially adept at encoding and integrating global visual information [53].

Pigeons are well known for their ability to process local information across various tasks [54–56]. These findings are very likely due to their ecology as granivorous birds that must discriminate small grains from a cluttered background using minute featural differences within their frontal visual field. The frontal field of pigeons is represented in the tectofugal pathway corresponding to the mammalian extrageniculocortical system [57, 58]. This system is far less prone to utilize global visual information than areas of the primate ventral cortical stream [46, 59]. This is reflected in the phenotypes, which show stronger reliance on pixel- and letter-level orthographic processing. Still, a cautious note is warranted, as the sequence-level representation shows relatively low reliability, making this result hard to interpret. Thus, neural adaptation to a granivorous lifestyle drives local precedence in pigeons and may potentially underlie their preference for pixel- and letter-level orthographic processing.

The differential specialization within a global-to-local processing continuum is not fixed in any of the three species we discuss. All of them have the remarkable ability to alter their search strategy in task- or training-dependent manners [60–62]. As we have observed in our analysis, this adaptability is a fascinating aspect of their cognitive strategies. While, on average, humans, baboons, and pigeons are positioned on a continuum of letter-sequence-, letter-, and pixel-level representation, each species shows considerable individual differences and is, in principle, capable of utilizing all three representations.

The finding of a substantial reliance on letter sequence representations in human orthographic decisions aligns with the general notion that efficient readers implement large orthographic elements for word recognition and reading (i.e., letter combinations or words; Ref. [2, 3, 63]). In literacy acquisition, increased reading competence strongly decreases the influence of the number of letters on reading time (i.e., longer reading times for longer words; e.g., Ref. [15, 64, 65]) and increases the perceptual span [5, 66, 67]. Still, when recognizing unknown letter strings (e.g., pseudowords), they rely again on letter units [68, 69]. The findings of this study show that, after training, human readers make orthographic decisions about non-words based on larger units that combine multiple letters in a sequence.

However, the implemented model variants fall short of simulating human behavior as accurately as the baboon and pigeon behavior. A potential explanation is that humans are the only species with previous experience with reading. We learned that the SLR can explain a high amount of variance based on visual-orthographic processing assumptions. Still, there is unexplained variance, which provides a clear motivation for further model development by integrating additional levels of representation into the computational model. Likely, candidate representations could be easily added to the model (while respecting their nature as prediction errors) and implemented at the orthographic, lexico-semantic, and phonological processing levels (e.g., phoneme-level prediction-error representations; see Appendix E). Here, human participants were efficient readers who learned that letter strings are associated with sound and meaning. However, we only modeled data from a pseudoword learning task (i.e., unknown letter strings; Ref. [23]), so it is likely that phonology played a role, given compelling evidence for its influence on visual word recognition (e.g., when learning to read; Ref. [15, 70]; in adult readers [6, 71–73]; when reading problems occur; Ref. [14, 74]). Another aspect is that, unlike more classical measures of orthography, the sequence-based prediction error implemented here respects the exact sequence, always starting with the first letter of the string. Here, we chose this representation as the first hypothesis at the sequence level, motivated by a word-length effect for pseudowords in adult reading [68, 69], which indicates sequential left-to-right letter-by-letter processing. However, the letter-sequence level implementation is only one of many potential representations that can characterize the orthographic structure of letter strings (e.g., representations based on bi- or tri-grams could have higher flexibility to model transposed letter effects; e.g., see Ref. [6, 18, 63]).

A limitation of the present study is that all stimuli were of fixed length (four letters for pigeons and baboons, five letters for humans). This design choice was motivated by the goal of investigating foveal word recognition (i.e., in contrast to parafoveal recognition processes; see [75, 76]) under controlled conditions and enabling direct cross-species comparison. However, natural reading involves variable word lengths, and the current implementations of the letter-position and letter-sequence representations are not tied to fixed-length inputs. Still, the use of a fixed letter length in our learned/novel categorization experiment is not necessarily artificial since, in natural reading, word length is perceived before fixation(e.g., see [76–79]). Previously, we systematically investigated different letter lengths of four to eight letters in humans and found an oPE effect in the expected direction for all lengths, using both length-specific and length-unspecific predictions (i.e., that highly correlate; r = .97; see [21]). Extending investigations of letter- and letter-sequence-level representations to variable-length strings and to interactions between foveal and parafoveal processing constitutes an important challenge for future modeling work.

Attention is currently not part of the SLR. However, in an exploration for LPE representations in humans, we implemented the well-established finding that the initial letter, the last letter, and the letter at the position of the fixation cross can be remembered and recognized better when compared to the letters in-between (e.g., [80–82]; see results in Appendix E). We find that attention-modulated LPE representations result in a larger difference between learned and novel items and greater variance explained when used in statistical models of human behavior. Future work is needed to establish potential differences in attention distributions between species. Yet, the present work demonstrates how computational models can be used to investigate systematically and in a qualitatively explicit manner the nature of representations involved in orthographic and other types of decisions, thereby also offering new possibilities to examine differences between individuals and species.

On a more general note, the Speechless Reader can be situated alongside other computational models of orthographic processing, including both cognitive and computer vision approaches (e.g., [42]). In particular, Agrawal et al. [83] propose a compositional model of lexical access based on visual letter similarity, operating at letter, bigram, and word levels. While this hierarchical structure superficially resembles the representational levels implemented in the Speechless Reader, the two models address different computational problems. The Speechless Reader is designed to account for lexical categorization (learned vs. novel strings) using predictive coding principles, whereas the model of Agrawal et al. aims to explain lexical access to specific word representations. As a consequence, direct model comparison is not straightforward in the context of the present task and research question, particularly given that word frequency and item familiarity were experimentally controlled in the current learning paradigm. Given that the current comparative study targets a categorization task across species rather than lexical access, we cannot directly compare the performance of our model to models of lexical access, even though similar representations are used.

## Conclusion

Our analyses reveal that humans, baboons, and pigeons implement orthographic decisions using different visuo-cognitive strategies. While human readers mainly relied on letter-sequence representations, baboons used letter- and letter-sequence-level strategies, whereas pigeons relied on all representations. These findings highlight the need for a “signature testing” approach that reconstructs how a task is solved [84]. Such an approach can uncover species-specific visuo-cognitive strategies and reveal the potential neuroevolutionary adaptations that drive these cognitive strategies. In the present study, this approach contributes to understanding the evolutionary precursors of orthographic processing in visual word recognition.

## Materials and methods

### Dataset descriptions: Participants, paradigm and stimuli

The data used for model simulations stems from Eisenhauer et al. [23] (Humans), Grainger et al. [8] (Baboons), and Scarf et al. [9] (Pigeons). Eisenhauer et al. [23] tested 37 human participants on a pseudoword lexical decision task of 880 trials, including eight presentations of 60 to-be-learned pseudowords (i.e., two presentations in each of the four sessions). Grainger et al. [8] used operant conditioning setups to train six baboons (*Papio papio*) in a word/non-word categorization task. Scarf et al. [9] trained four pigeons (*Columba livia*) on a similar task. To investigate the phenotype of the trained participants across species, we limited our primary analysis to the final 10,000 trials for each baboon and pigeon, and the last two sessions in the human dataset, for a total of 440 trials per participant. Furthermore, the limitation of the learned items led to an adaptation of the lexical process part of the model [6], allowing the model to be adapted to individual learner thresholds for each participant (see below).

All three datasets were, in principle, based on a two-forced-choice task, without time pressure, that indicated whether a letter string was learned or not [8, 9, 23]. Still, task implementation differed in several parameters that were, to the main objective of the task, rather cosmetic: Humans received black stimulation on a white background, starting with fixation bars, followed by the presentation of a letter string in Courier New font until a button press. Baboons and pigeons, in contrast, received yellow letter-string stimulation on a black background in Arial font, combined with binary-choice touchscreen indicators. Learned and novel letter strings consisted of 5 letters in the human task and 4 letters in the animal task. The most relevant word characteristics (i.e., the prediction-error representations) depend on the individual lexicons inferred from individual performance (see Supplemental Table 1). So these word characteristics differ for each individual and are, therefore, a first result, described in the results section. For further information on participants, analysis, and stimulus materials, see this OSF repository (here).

### Model implementation

Instead of automatically estimating the categorization boundary (blue line in Fig. 1) based on the overlap of word-likeness estimates from words and non-words, we implemented the categorization process based only on the word-likeness distribution of the learned letter strings (also described in Ref. [6]). To find an adequate categorization threshold, we calculated the model responses for all possible model variants and multiple possible threshold values (range 5 - 95; see Eq. 6) for each participant (see Supporting Information: Table S1 and Figs. S1, S2, S3). To determine the optimal boundary, we compared the model simulations to the behavior of each participant.

In all of our model simulations, we investigate behavior after initial training (i.e., when animal performance is relatively stable). For all datasets, the items in the lexicon are determined based on performance in the trials used in the analysis (all trials after the first 33,041, i.e., determined by the available data for both baboons and pigeons, and by data from sessions 3 and 4 of the human dataset). We use the criterion for learned items, which was previously reported (Accuracy *>* .71), to determine the participant-specific implementation of the items in the individual lexicons of the speechless reader. When the lexicon items are known, we can estimate the top-down predictions (gray lines in Figure 1) on the level of pixels, letters, and letter sequences (see Eq. 1, described in Ref. [20, 21, 30]). When estimated on the pixel level, the *StoredItems* matrix consists of gray values defining the image of each item stored in the lexicon (*i*,*j* = 140, 40 each). For letters, the *StoredItems* matrix consists of all letters (for four-letter words: *i*,*j* = 4, 26 each); and for the letter sequence level, of all possible letter sequences (e.g., for four-letter words: Letters 1 and 2; Letters 1, 2, and 3; *i*,*j* = 2, x where x is the number of all possible letter combinations for the two and three letter sequences). For each pixel, letter, and letter sequence, a mean over the number of items in the lexicon (*n*) is calculated to represent the prediction. Note that letter-based representations were informative. In the animal studies, the letter frequencies ranged from 2.6 to 5.3% at each of the five positions. Also, explorations with inverted sequence prediction errors (i.e., starting the sequence with the last letter of a word) yielded much lower model simulation accuracy, indicating that animals used sequence representations in the reading direction.

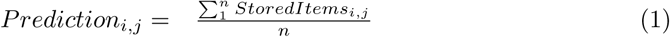

Next, we integrate the predictions with the sensory input by subtraction (see Formula 2). The *SensoryInput* matrix is similar to the prediction on each level, so the subtraction can be implemented for each letter or letter sequence. Note that we represent the sensory input as 1 at the letter and letter-sequence levels, indicating the maximum possible error when the input is unpredictable. This value is reduced by the prediction value for the letter at the given position (e.g., in English, there is a high probability for the *s* being the last letter, so we expect a low prediction error at this position) or for the letter sequence relevant (i.e., *th* is likely for English for the initial letter sequence). We further developed the formulation for the pixel level (*oPE*). We conjectured that a binary prediction error would be most appropriate when modeling word/non-word categorization behavior (see Eq. 3 and [20]). We found that a threshold of 0.5 is well-suited to implement the binary prediction error representation. After integrating prediction and sensory input to generate the prediction error, all matrix values are summed to obtain a single value summarizing the overall prediction error at each prediction error level for each letter string.

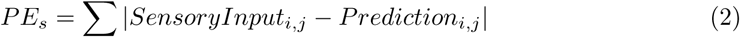

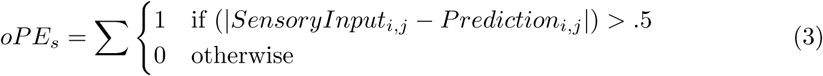

We calculated prediction errors at all three levels for each learned and novel letter string in the datasets. In preparation for the integration of multiple prediction errors to estimate word-likeness based on prediction errors, we normalized each set of prediction errors to a value between 0 and 1 (*PE_norm_*; see Eq. 10). After that, we can accumulate multiple prediction error values by summation and division (see Eq. 5). The resulting prediction error estimate is normalized across all model variants that integrate multiple prediction error representations.

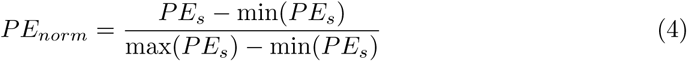

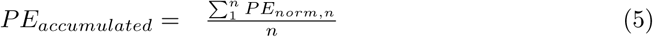

In the final step, we implement the lexical categorization process. Initially, we implemented the boundary at the highest lexical categorization difficulty (i.e., when the distributions of prediction-error-based word-likeness for learned and novel letter strings overlapped most, see [6]). Here, we conceived of an alternative approach, with a similar logic, that would allow us to implement lexical categorization without including novel letter strings. This is potentially more realistic, as the storage of non-words is unlikely, and it works better when the number of items in the lexicon is low, which is the case for some individuals. We achieved this by considering multiple thresholds applied to the prediction errors of the learned letter strings (see Eq. 6). Since we could not automatically calculate the optimal threshold, which would have been the case in the original formulation [6], multiple thresholds in the range from 5 − 95 were applied to all model variants to achieve a binary value, which indicates whether a letter string is learned or not (see the blue line in Fig. 1). Interestingly, this alternative approach resulted in better model fits than the original implementation used for baboons in the past (i.e., cp. Ref. [30]). Thus, for the present analysis, we focused on the approach that can be implemented without assuming that non-words must be stored.

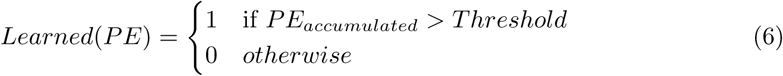

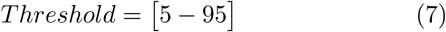

With this model, we simulated the behavior of each participant with each model variant (i.e., all prediction-error combinations with all possible thresholds; N = 7). To determine the most appropriate model, we compared model and participant behavior across all trials using the mean squared error metric (see Fig. S1-3a; see also mean model accuracy in Fig. S1-3b). We selected the threshold value that yielded the smallest absolute difference between the data and simulations across all model variants. Note that a different lexicon is assumed for each participant, which is why simulations differed between participants. The model variant with the lowest mean squared error was considered to include the prediction error representations most likely implemented to achieve the orthographic decision performance of the individual participant (see Figure 3c,f,i and Table S1). The model was implemented in Python, and the scripts are available here (find here).

### Data analysis

The description of the statistical analysis has three parts: (i) Analysis of model parameters, (ii) analysis of response data, and (iii) analysis of model fit and stability.

For the analysis of model parameters, the differentiation of the prediction error representations between learned and novel items is of main interest. To test whether a representation differs between the two types of stimuli, we ran a linear mixed model of the following form:

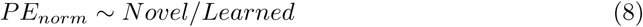

using the *lme*4 package implemented in GNU R [85]. For the random-effect estimates, we included random intercepts for participants and letter strings in all models. We estimated the random slope of Novel vs. Learned for participants when estimating the oPE effect, following the concept of parsimonious mixed models [86]. Furthermore, we estimated *p*-values with the *lmeTest* package using the Satterthwaite approximation [86] and corrected the *p*-level for multiple comparisons using a Bonferroni correction, leading to a *p*-level of 0.016 (three tests per prediction error for each species; [87, 88]).

To measure the effect of the prediction error representations on behavior, we used generalized linear mixed model analysis of accuracy data (presented in the results section) and linear mixed model analyses and response time comparisons (presented in the Appendix A, Table S1, Figure S1). For the accuracy analysis, we modeled the raw binary outcome measure using a binomial distribution with the *glmer*() function from Bates et al. [85], again implementing parsimonious mixed models and the Bonferroni correction. Modelling binomial data is highly susceptible to increased model complexity, which can lead to non-convergence. Thus, for accuracy, we modelled each of the three levels of representation separately and estimated random effects on the intercept only (see [86]). In contrast, for the analysis of response times, we combined all three interaction terms into a single model, as the linear mixed model for interval-scaled data can handle much more complex models (i.e., via the *lmer*() function), including random-slope estimation. Hence, we could model them at once. Note that, since response time distributions are typically ex-Gaussian, we transformed the response times using a natural logarithm to achieve normally distributed data. We used the following formula

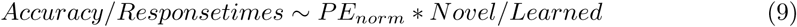

using random effect estimates for the intercept of both individuals and Letter strings and estimating all possible random slopes for the Individuals’ random effect, including the interaction term.

To estimate model fit and stability, we first tested the simulation’s fit to the data using a linear regression model with the *lm*() function in base R. With the following formula

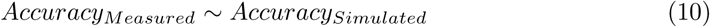

We are investigating the similarity of individual differences between model simulations and participant behavior. For this analysis, we aggregated the data to the individual level using an arithmetic mean for each species. Thus, for each individual, we had one measured and simulated mean accuracy for each stimulus condition. In addition, to investigate the group-level similarity, we provide descriptive statistics, including boot-strapped 95% confidence intervals.

To examine the stability of the implemented prediction error representations, we conducted 100 split-half reliability estimations per individual (e.g., see [29] for the importance of reliability estimates in the context of computational phenotyping). We estimated each individual’s phenotype twice, once for each data half. After that, we compared the two phenotypes of the same individual and estimated the overlap in the representations of the models fitted to either of the two halves. The resulting metric is the mean overlap of representations across the two splits, expressed as a percentage (i.e., 100% indicates that the phenotypes were present in all iterations and splits).

## Acknowledgments

We thank Ulrike Basten for helping with the reliability estimation. This research was funded by the German Research Foundation (Grant no. 523332674 awarded to B.G.), the European Research Foundation (Project AVIAN MIND ERC-2020-ADG, LS5, GA No. 101021354 awarded to O.G.; FP7/2013, Grant 617891 awarded to C.J.F.), and the Royal Society of New Zealand (Marsden Fund grant 19-UOO-162 awarded to M.C.).

## Appendix A

**Fig S1.**
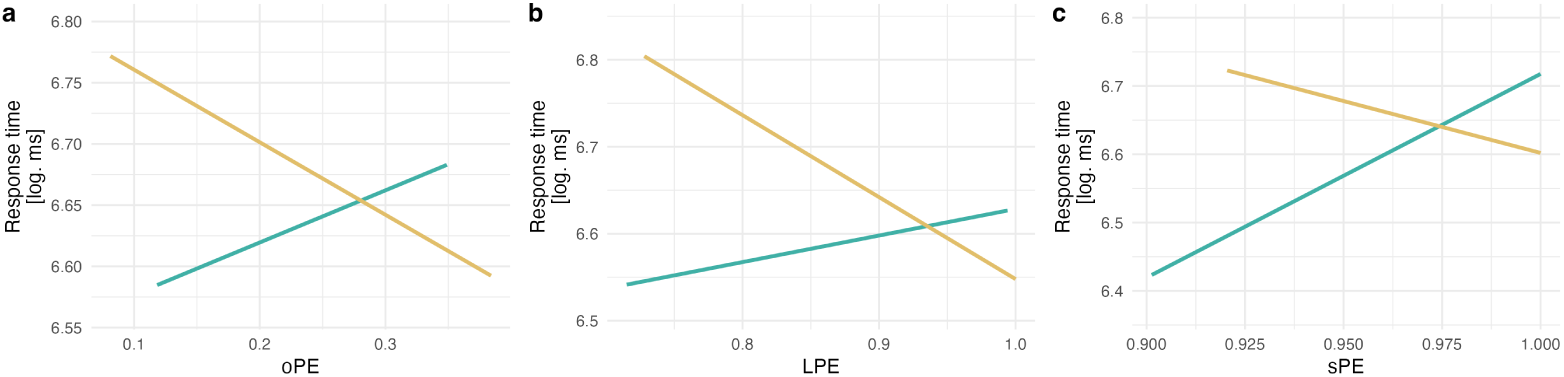
(a) oPE, (b) LPE, and (c) sPE effects on decision response times with dark yellow showing the effect on novel and green showing the effect on learned letter strings. See the Table below for the statistics.

**Table S1.** Statistics of the linear mixed model with response times as the dependent variable, testing the interaction of the three prediction errors with stimulus condition (Learned vs. Novel).

|  | Estimate | SE | t-value |
| --- | --- | --- | --- |
| **Intercept** | 5.3 | 0.4 | 13.9 |
| **Main effects** |  |  |  |
| Learned/Novel | 4.8 | 0.5 | 8.9 |
| oPE | 0.3 | 0.3 | 1.2 |
| LPE | 0.4 | 0.3 | 1.5 |
| sPE | 1.0 | 0.4 | 2.2 |
| **Interactions** |  |  |  |
| oPE X Learned/Novel | -0.6 | 0.3 | -2.1 |
| LPE X Learned/Novel | -1.2 | 0.3 | -3.7 |
| sPE X Learned/Novel | -3.6 | 0.6 | -6.0 |

## Appendix B

**Fig S2.**
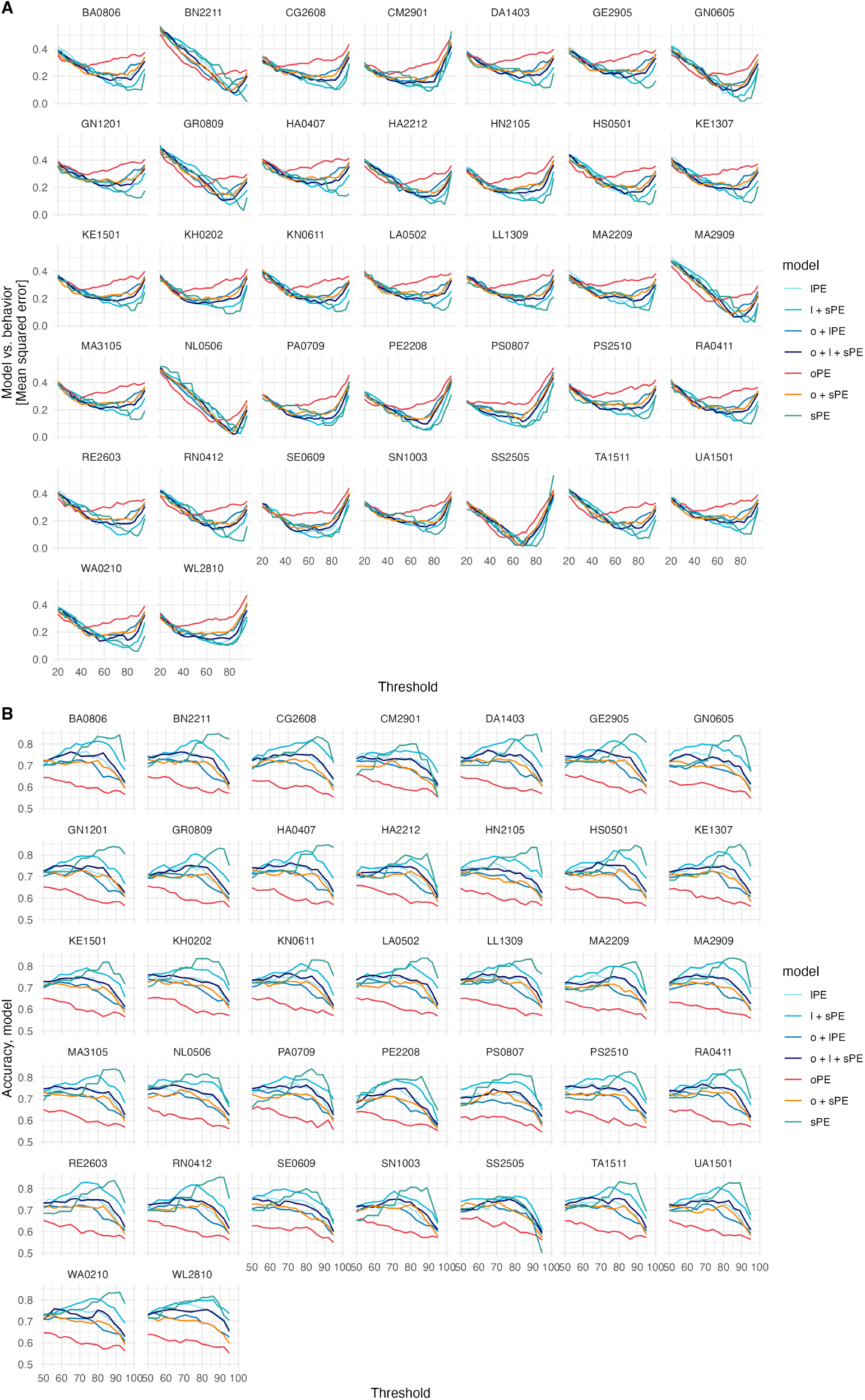
Model comparisons and model accuracy for all models and all humans. (A) Shows the mean squared error of each model (i.e., all combinations of representations and all tested thresholds) and human behavior. (B) Model accuracy from all tested models.

**Fig S3.**
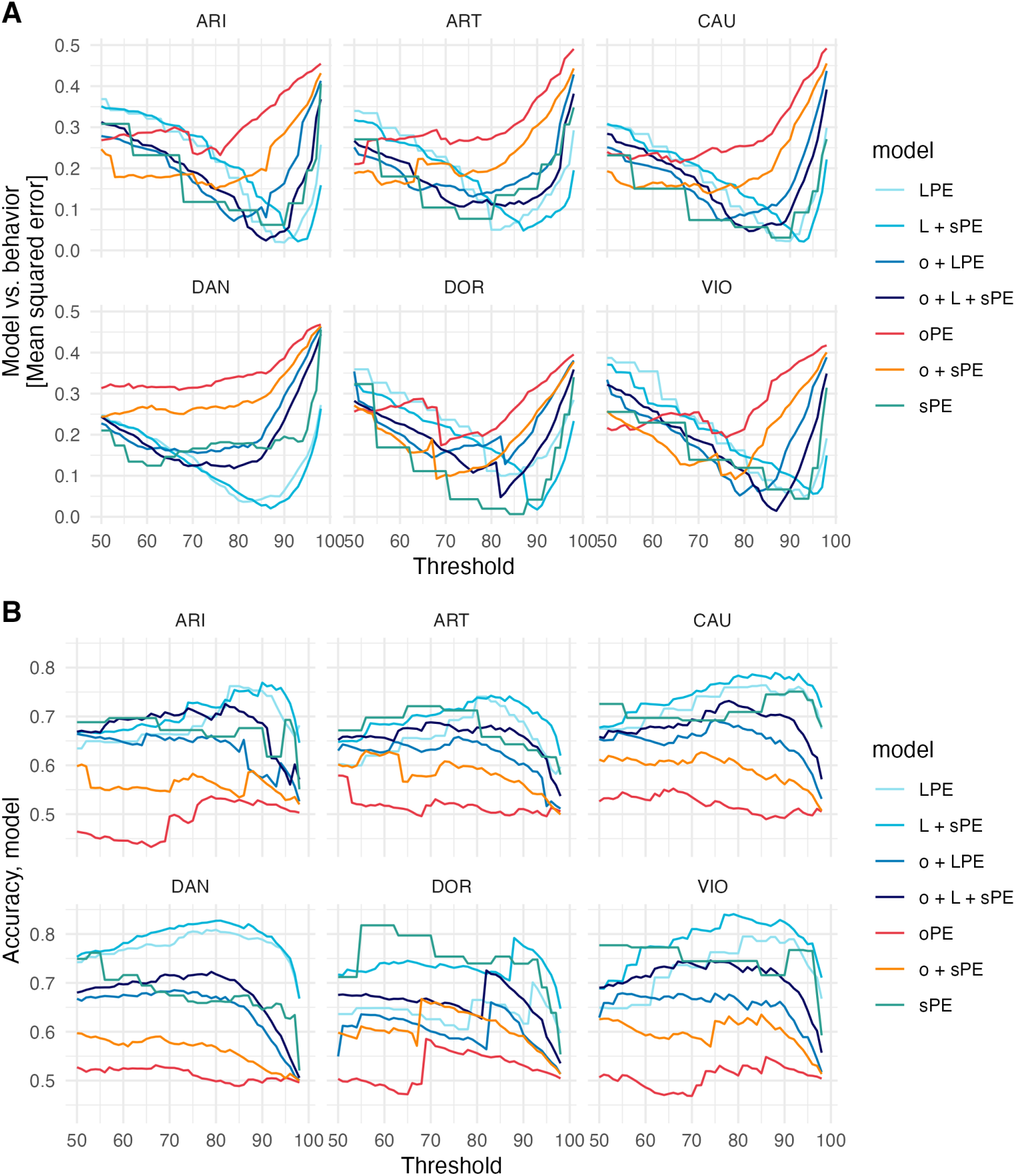
Model comparisons and model accuracy for all models and all humans. (A) Shows the mean squared error of each model (i.e., all combinations of representations and all tested thresholds) and baboon behavior. (B) Model accuracy from all tested models.

**Fig S4.**
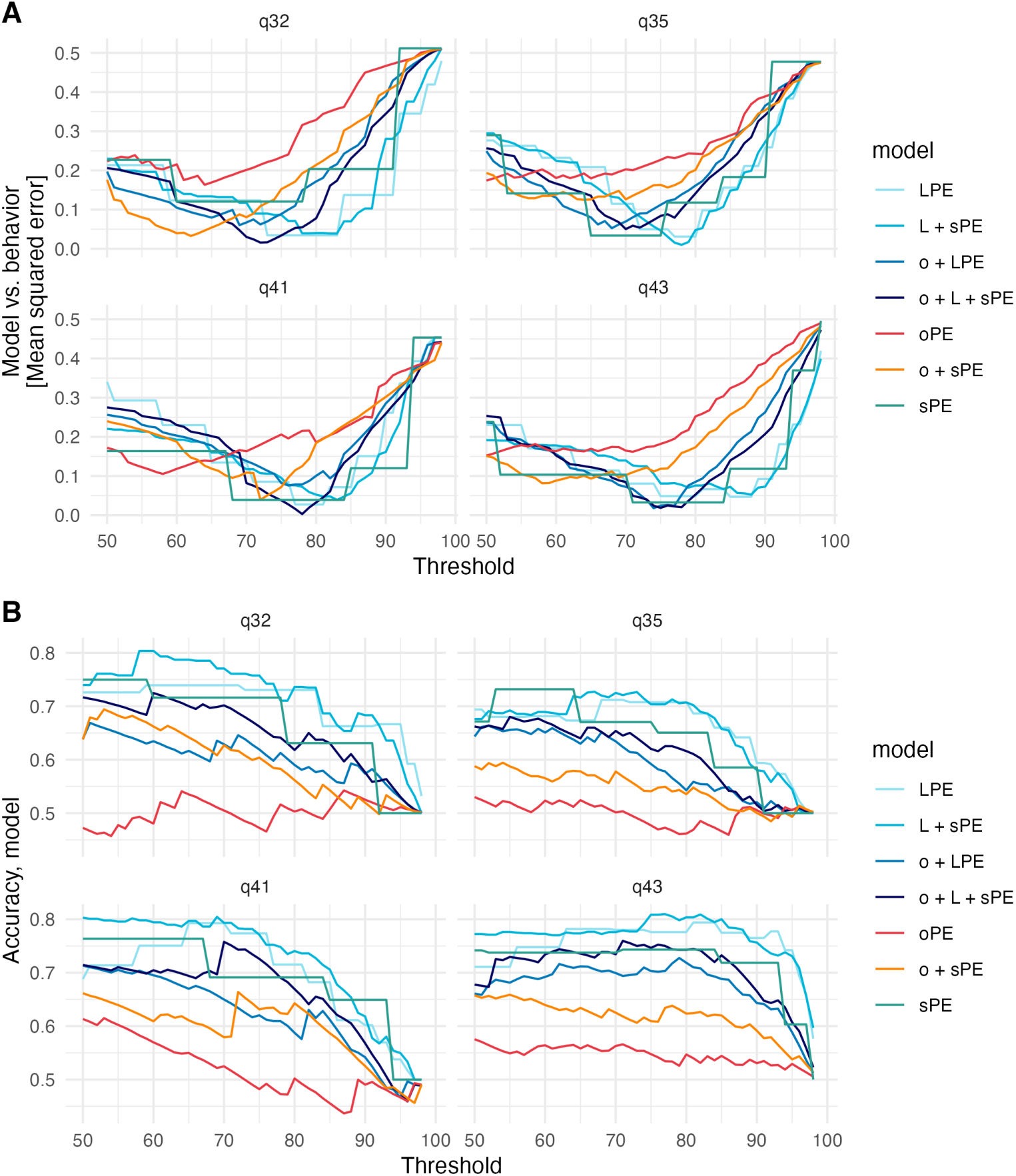
Model comparisons and accuracy for all models and all pigeons. (A) Shows the mean squared error of each model (i.e., all combinations of representations and all tested thresholds) and pigeon behavior. (B) Model accuracy from all tested models.

**Fig S5.**
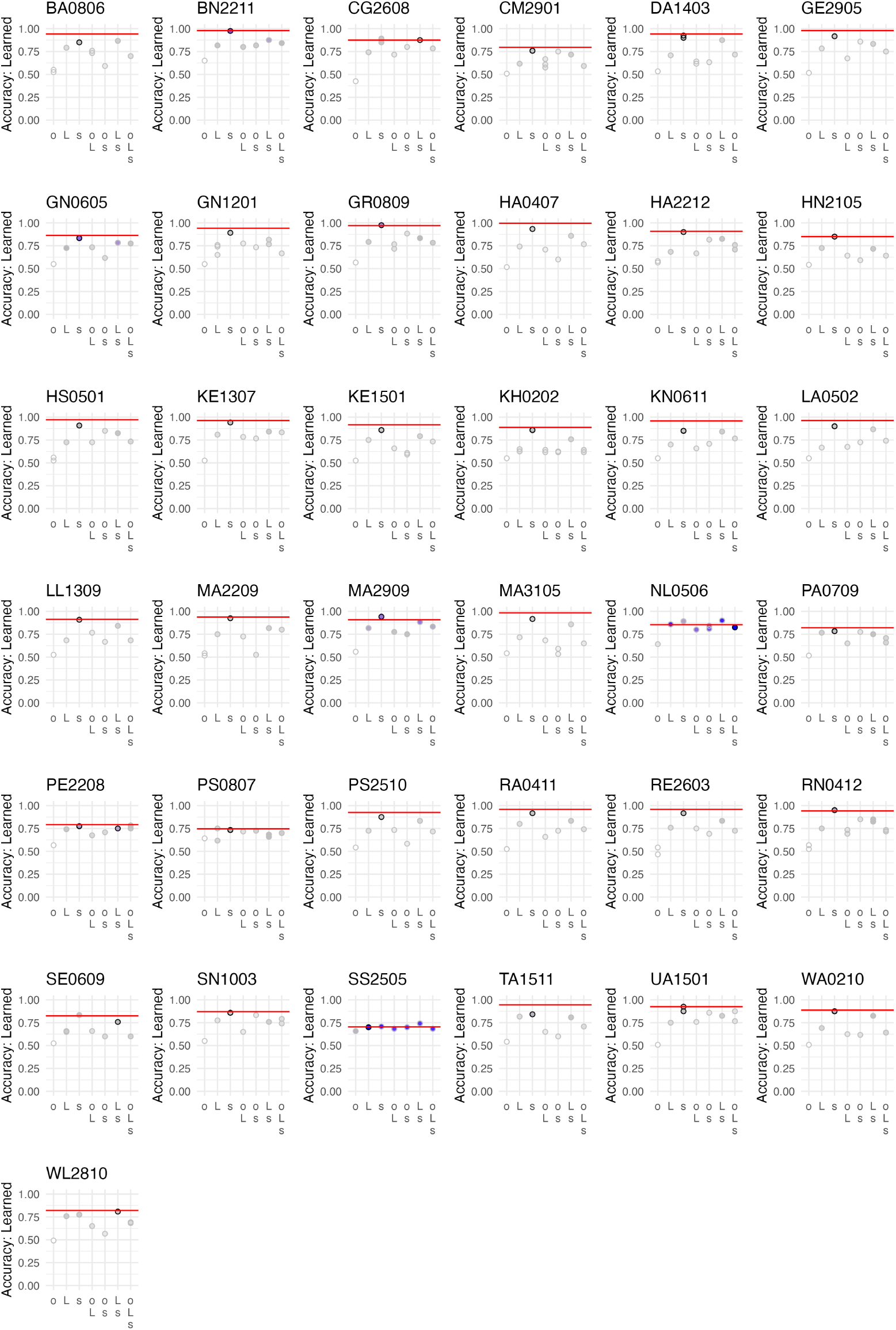
Human individual error analysis of the accuracy of learned items. Dots represent the best-fitting models from each phenotype indicated in the y-axis labels. Black rings indicate the individual’s phenotype, dot color indicates the mean squared error, darker colors indicate the lowest values, and white indicates the highest values. The red line indicates the individual’s performance.

## Appendix C

**Table S2.** Number of words in the lexicon for each participant. Also, mean squared error, representations, and thresholds for the winning model for each participant.

|  | N | Error | oPE | LPE | sPE | Thres. |
| --- | --- | --- | --- | --- | --- | --- |
| Human/1 | 56 | 0.10 | 0 | 0 | 1 | 80 |
| Human/2 | 59 | 0.01 | 0 | 0 | 1 | 95 |
| Human/3 | 49 | 0.10 | 0 | 1 | 1 | 83 |
| Human/4 | 45 | 0.09 | 0 | 0 | 1 | 77 |
| Human/5 | 55 | 0.12 | 0 | 0 | 1 | 89 |
| Human/6 | 59 | 0.11 | 0 | 0 | 1 | 89 |
| Human/7 | 51 | 0.02 | 0 | 0 | 1 | 83 |
| Human/8 | 53 | 0.12 | 0 | 0 | 1 | 89 |
| Human/9 | 58 | 0.03 | 0 | 0 | 1 | 92 |
| Human/10 | 59 | 0.14 | 0 | 0 | 1 | 92 |
| Human/11 | 55 | 0.08 | 0 | 0 | 1 | 89 |
| Human/12 | 48 | 0.09 | 0 | 0 | 1 | 86 |
| Human/13 | 57 | 0.07 | 0 | 0 | 1 | 89 |
| Human/14 | 59 | 0.08 | 0 | 0 | 1 | 89 |
| Human/15 | 58 | 0.11 | 0 | 0 | 1 | 86 |
| Human/16 | 52 | 0.09 | 0 | 0 | 1 | 83 |
| Human/17 | 55 | 0.11 | 0 | 0 | 1 | 86 |
| Human/18 | 56 | 0.12 | 0 | 0 | 1 | 89 |
| Human/19 | 55 | 0.11 | 0 | 0 | 1 | 89 |
| Human/20 | 55 | 0.11 | 0 | 0 | 1 | 89 |
| Human/21 | 57 | 0.03 | 0 | 0 | 1 | 92 |
| Human/22 | 58 | 0.13 | 0 | 0 | 1 | 89 |
| Human/23 | 42 | 0.02 | 1 | 1 | 1 | 83 |
| Human/24 | 44 | 0.09 | 0 | 0 | 1 | 80 |
| Human/25 | 50 | 0.05 | 0 | 0 | 1 | 77 |
| Human/26 | 40 | 0.07 | 0 | 0 | 1 | 71 |
| Human/27 | 56 | 0.13 | 0 | 0 | 1 | 86 |
| Human/28 | 54 | 0.10 | 0 | 0 | 1 | 89 |
| Human/29 | 54 | 0.09 | 0 | 0 | 1 | 89 |
| Human/30 | 54 | 0.05 | 0 | 0 | 1 | 92 |
| Human/31 | 50 | 0.08 | 0 | 1 | 1 | 68 |
| Human/32 | 52 | 0.10 | 0 | 0 | 1 | 83 |
| Human/33 | 34 | 0.01 | 0 | 1 | 0 | 68 |
| Human/34 | 56 | 0.07 | 0 | 0 | 1 | 80 |
| Human/35 | 52 | 0.10 | 0 | 0 | 1 | 89 |
| Human/36 | 52 | 0.06 | 0 | 0 | 1 | 89 |
| Human/37 | 48 | 0.10 | 0 | 1 | 1 | 77 |
| Baboon/ARI | 67 | 0.04 | 1 | 1 | 1 | 86 |
| Baboon/ART | 87 | 0.11 | 0 | 1 | 1 | 88 |
| Baboon/CAU | 73 | 0.04 | 0 | 1 | 1 | 93 |
| Baboon/DAN | 271 | 0.03 | 0 | 1 | 1 | 87 |
| Baboon/DOR | 96 | 0.04 | 0 | 0 | 1 | 87 |
| Baboon/VIO | 86 | 0.03 | 1 | 1 | 1 | 87 |
| Pigeon/Q32 | 8 | 0.01 | 0 | 1 | 0 | 73 |
| Pigeon/Q35 | 28 | 0.03 | 0 | 1 | 1 | 75 |
| Pigeon/Q41 | 21 | 0.05 | 1 | 1 | 0 | 77 |
| Pigeon/Q43 | 32 | 0.04 | 1 | 1 | 1 | 75 |

## Appendix D: Individual error analysis

**Fig S6.**
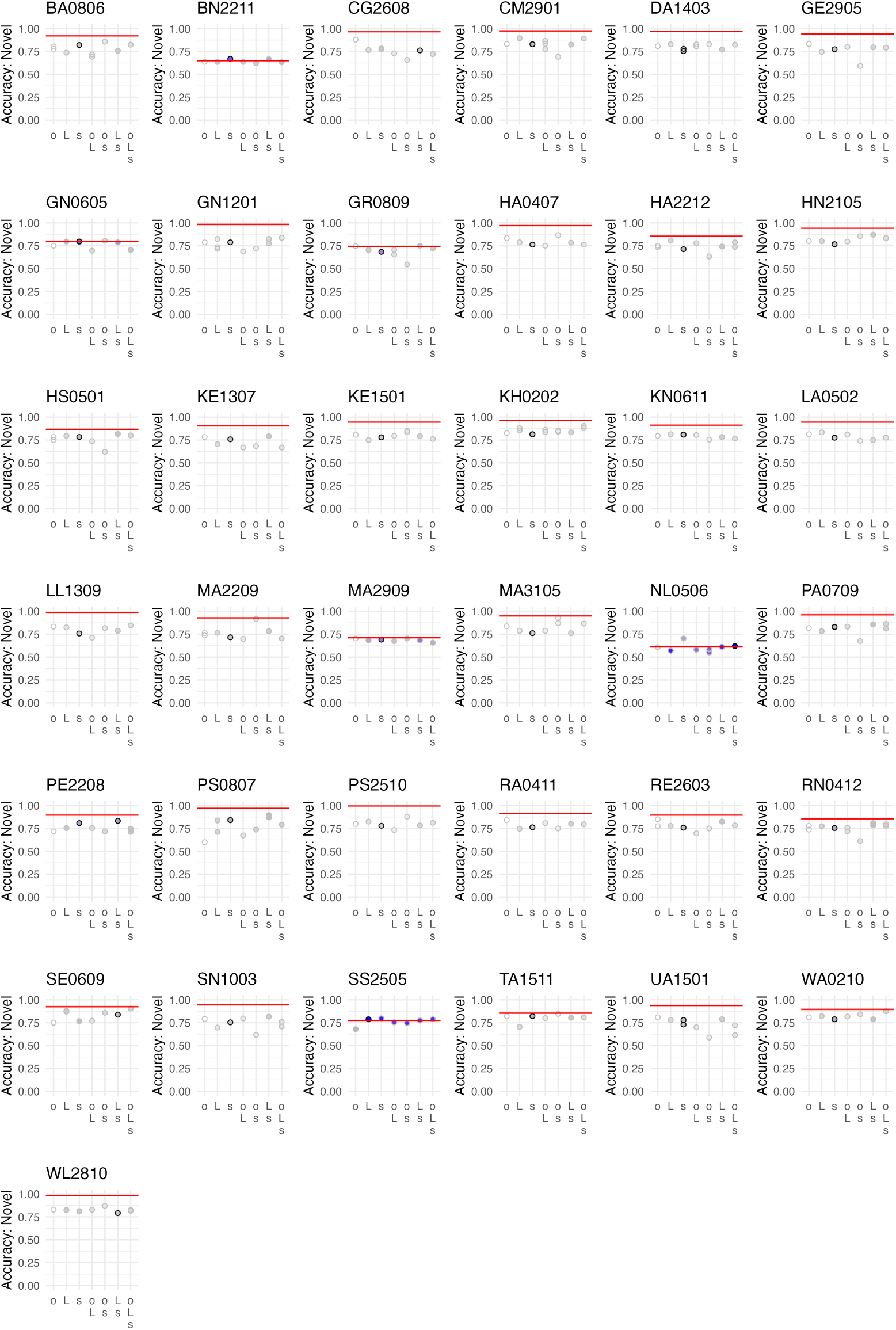
Human individual error analysis of the accuracy of novel items. Dots represent the best-fitting models from each phenotype indicated in the y-axis labels. Black rings indicate the individual’s phenotype, dot color indicates the mean squared error, darker colors indicate the lowest values, and white indicates the highest values. The red line indicates the individual’s performance.

**Fig S7.**
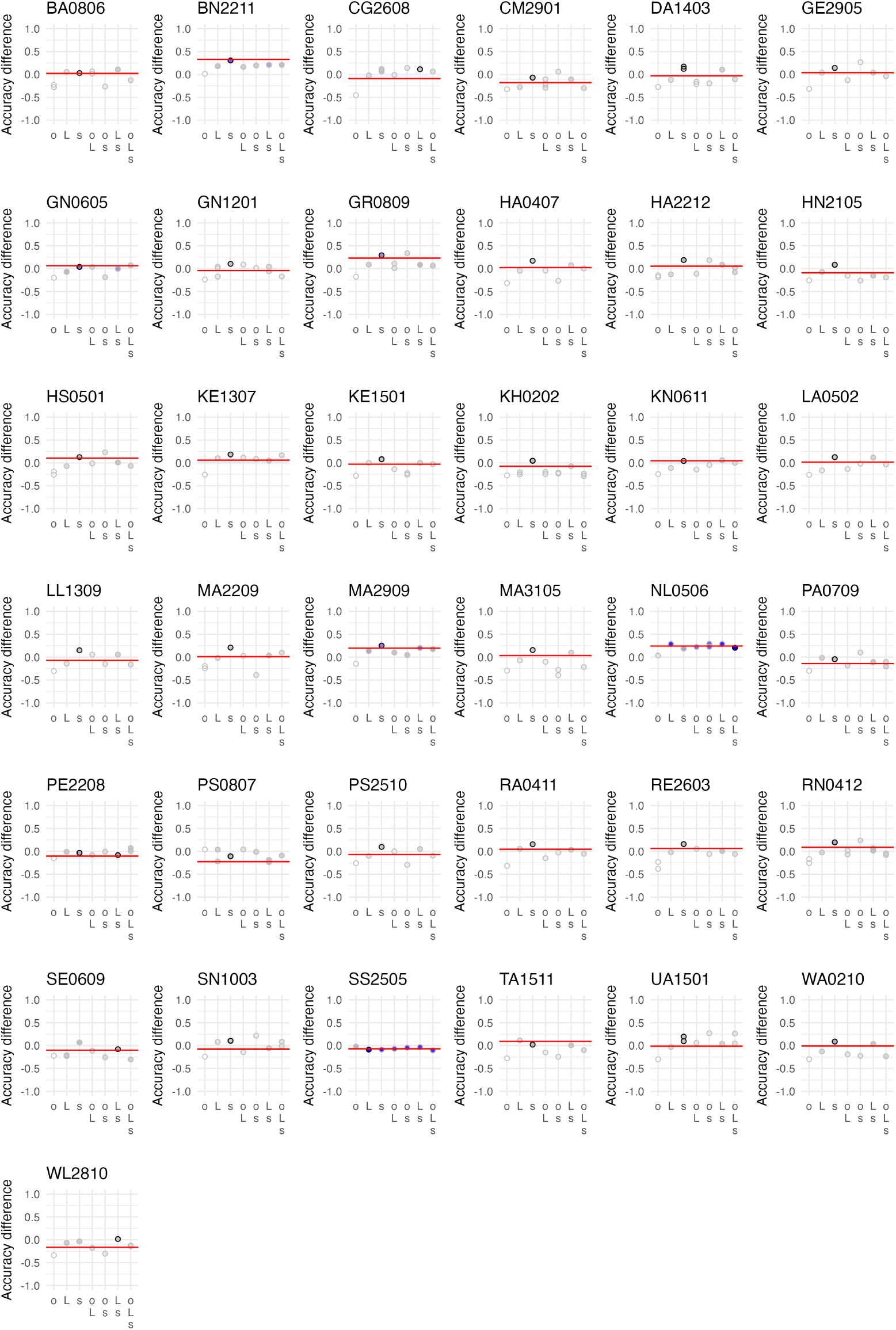
Human individual error analysis of the accuracy difference between learned and novel items. Dots represent the best-fitting models from each phenotype indicated in the y-axis labels. Black rings indicate the individual’s phenotype, dot color indicates the mean squared error, darker colors indicate the lowest values, and white indicates the highest values. The red line indicates the individual’s performance.

**Fig S8.**
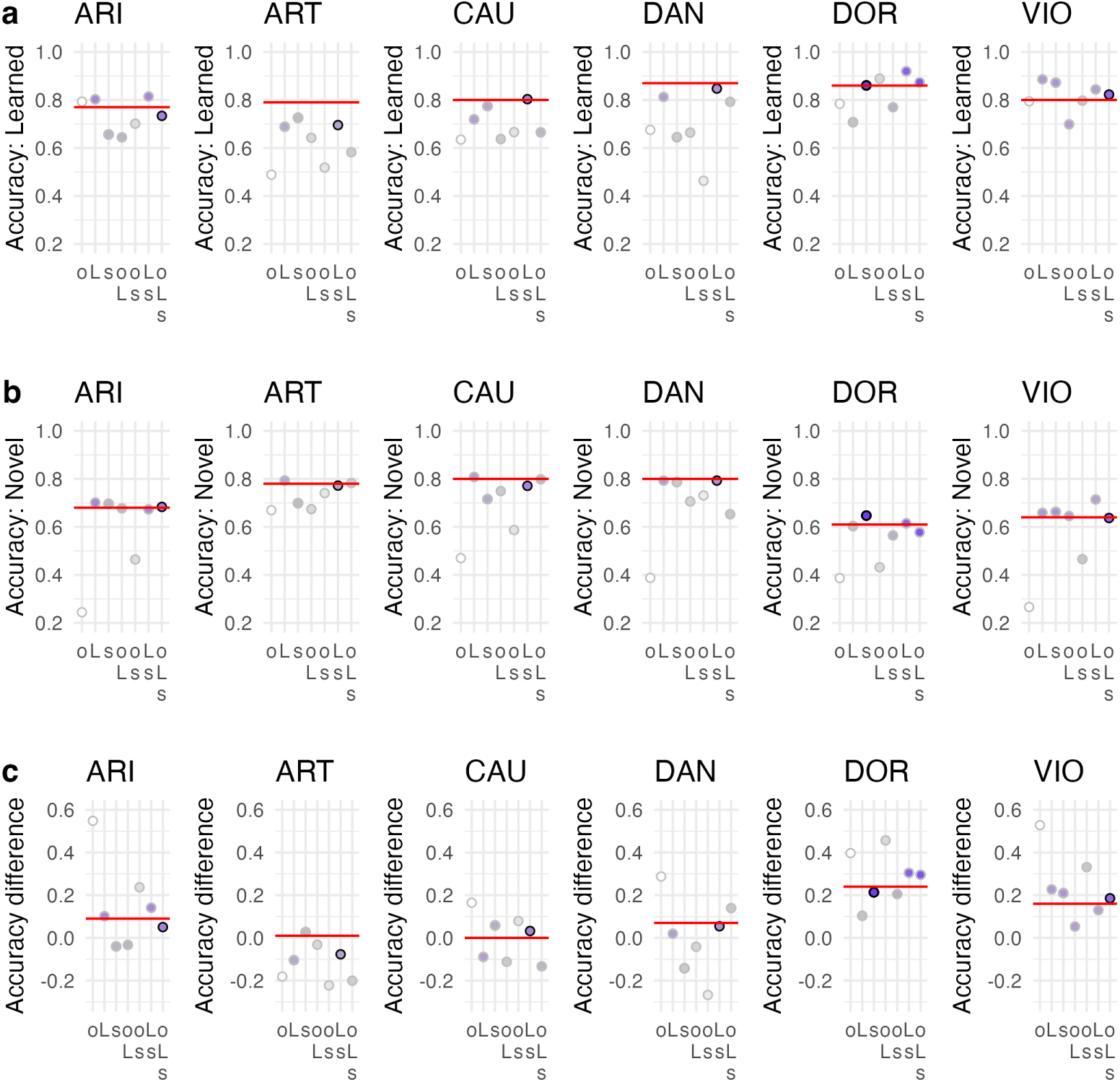
Individual error analysis for all four baboons. Top row: Accuracy for learned items; Middle row: Accuracy for novel items; Lower row: Accuracy difference between learned and novel items. Dots represent the best-fitting models from each phenotype indicated in the y-axis labels. Black rings indicate the individual’s phenotype, dot color indicates the mean squared error, darker colors indicate the lowest values, and white indicates the highest values. The red line indicates the individual’s performance.

**Fig S9.**
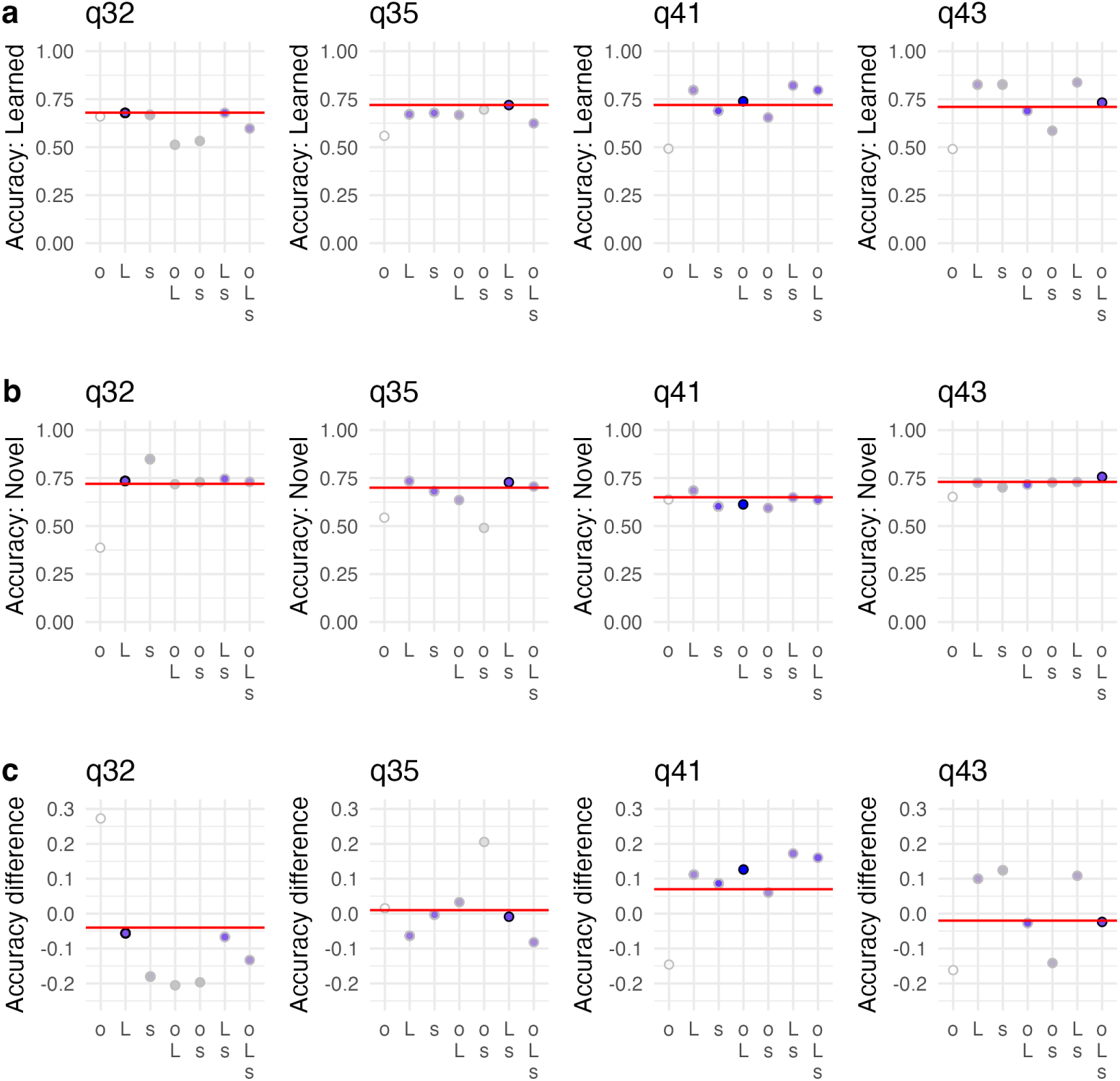
Individual error analysis for all four pigeons. Top row: Accuracy for learned items; Middle row: Accuracy for novel items; Lower row: Accuracy difference between learned and novel items. Dots represent the best-fitting models from each phenotype indicated in the y-axis labels. Black rings indicate the individual’s phenotype, dot color indicates the mean squared error, darker colors indicate the lowest values, and white indicates the highest values. The red line indicates the individual’s performance.

## Appendix E: Implementation of letter-based attention for humans

Here, we explore the possibility of implementing letter-level attention to the speechless reader for humans. Evidence from human readers typically shows that attention follows a pattern that favors both the initial and the final letter of a word as well as the central letter [81–83]. For this initial exploration, we focus on letter-level prediction-error representations, for which the implementation is straightforward. Such that, after the letter level prediction errors are established for each letter, we multiply the prediction error matrix by an attention matrix (i.e., multiplying the single letter LPE with a single letter attention value). For example, if the prediction error matrix is as follows [.7*, .*9*, .*9*, .*8*, .*9] and the attention matrix is as follows [1, .5, 1, .5, 1], the result from the muliplication would be a LPE matrix of [.7*, .*45*, .*9*, .*4*, .*9] (i.e., when inspecting the fourth letter one would multiply .8 with .5 resulting in an attention weighted LPE of .4). When implementing the expected attention reduction for the second and fourth letter, we compared model implementations that reduced attention by [.25*, .*5*, .*75*, .*85*, .*95, 1], with 1 representing the standard implementation. Thus, we can explore which attention-reduction assumption best predicts human responses.

First, we investigate how the differences in the LPE between novel and learned letter strings change when attention is applied. We find the highest difference between the LPE estimation when attention is lowest to the second and fourth letter (see Table S3). Thus, the differentiability between the letter strings increases with lower attention, indicating that differentiation between letter strings should be easier.

**Table S3.** Comparison of mean LPE values from novel and learned letter strings, including the difference comparing different attentional weights for the Letters 2 and 4. Note that an attention value of 1 indicates no implementation of attention, as all letters are weighted with the same value.

| Attention value | LPE: Novel | LPE: Learned | Difference |
| --- | --- | --- | --- |
| .25 | 0.72 | 0.60 | 0.12 |
| .50 | 0.78 | 0.69 | 0.10 |
| .75 | 0.85 | 0.77 | 0.08 |
| .85 | 0.87 | 0.80 | 0.07 |
| .95 | 0.89 | 0.83 | 0.06 |
| 1 | 0.91 | 0.85 | 0.06 |

When directly testing the attention assumptions on behavior (i.e., accuracy and response times), we use statistical model comparisons. We find that the LPE parameters, including attention, improved model fit across both accuracy and response times. For accuracy, the best model assumed an attentional reduction value of .85, and for response times, a value of .75 (i.e., showing the lowest AIC value; Fig. S4). This result shows that the increase in differentiability from attentional weighting also yields a better description of the behavioral data. Thus, this initial exploration demonstrates that the implementation of attention has the potential to implement a more realistic model for human visual word recognition behavior. Future work must treat attention as a fundamental algorithm that influences not only one level of the model but all levels, potentially starting at the visual level. Also, the implementation of attention should be parsimonious and theoretically driven, trying to implement a model with a minimal set of explicit assumptions, as the speechless reader is for animal behavior.

**Table S4.** Statistical model comparison investigating different attentional weights: AIC model fit statistics for Human accuracy and response times. The model in bold letters/numbers is the winning model, as indicated by the lowest AIC value, which corresponds to the highest model fit.

| Model | Attention value | N par. | AIC |
| --- | --- | --- | --- |
| **Accuracy** |  |  |  |
| H0 (no LPE included) | - | 4 | 10015.9 |
| Equal attention distribution | 1 | 6 | 8776.8 |
| Attention reduction for Letter 2 & 4 | .25 | 6 | 9254.0 |
| Attention reduction for Letter 2 & 4 | .50 | 6 | 9032.9 |
| Attention reduction for Letter 2 & 4 | .75 | 6 | 8770.0 |
| Attention reduction for Letter 2 & 4 | **.85** | 6 | **8697.6** |
| Attention reduction for Letter 2 & 4 | .95 | 6 | 8703.6 |
| **Response times** |  |  |  |
| H0 (no LPE included) | - | 5 | 13231.0 |
| Equal attention distribution | 1 | 7 | 13097.0 |
| Attention reduction for Letter 2 & 4 | .25 | 7 | 13111.0 |
| Attention reduction for Letter 2 & 4 | .50 | 7 | 13086.0 |
| Attention reduction for Letter 2 & 4 | **.75** | 7 | **13069.0** |
| Attention reduction for Letter 2 & 4 | .85 | 7 | 13072.0 |
| Attention reduction for Letter 2 & 4 | .95 | 7 | 13086.0 |

## Appendix F: Visualization aiding Table 5

**Fig S10.**
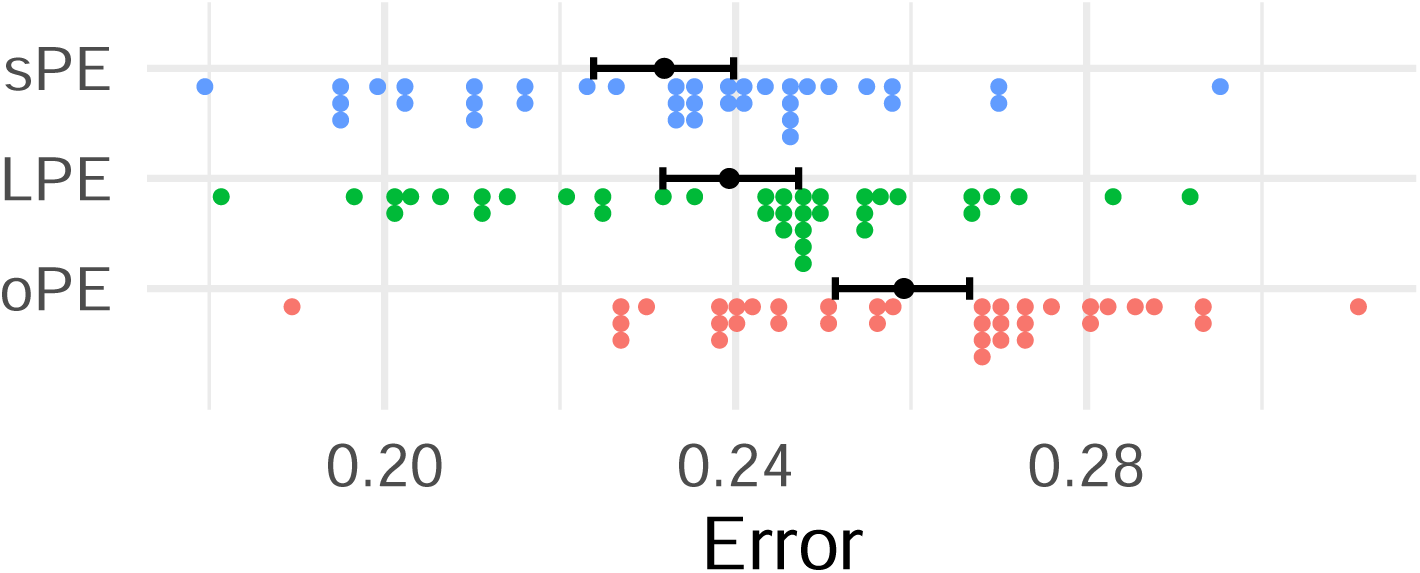
Model independent mean squared error, for each of the representations aggregated to participant level for each human. Colored dots represent individual errors, the black dot represents the mean, and whiskers represent the 95% CI.

## Notes

### Competing Interest Statement

The authors have declared no competing interest.

### Summary of Updates

- New Pigeon data leading to a adapted interpretation of findings - Additional analysis (e.g., error analysis, reliability analysis) - Extended methods section

## References

1. Dehaene S, Cohen L. The unique role of the visual word form area in reading. Trends in Cognitive Sciences. 2011;15(6):254–262. doi:10.1016/j.tics.2011.04.003.

2. Coltheart M, Rastle K, Perry C, Langdon R, Ziegler J. DRC: A dual route cascaded model of visual word recognition and reading aloud. Psychological Review. 2001;108(1):204–256. doi:10.1037/0033-295x.108.1.204.

3. Perry C, Ziegler JC, Zorzi M. Nested incremental modeling in the development of computational theories: The CDP model of reading aloud. Psychological Review. 2007;114(2):273–315. doi:10.1037/0033-295x.114.2.273.

4. Engbert R, Nuthmann A, Richter EM, Kliegl R. SWIFT: A Dynamical Model of Saccade Generation During Reading. Psychological Review. 2005;112(4):777–813. doi:10.1037/0033-295x.112.4.777.

5. Rayner K. The 35th Sir Frederick Bartlett Lecture: Eye movements and attention in reading, scene perception, and visual search. Quarterly Journal of Experimental Psychology. 2009;62(8):1457–1506. doi:10.1080/17470210902816461.

6. Gagl B, Richlan F, Ludersdorfer P, Sassenhagen J, Eisenhauer S, Gregorova K, et al. The lexical categorization model: A computational model of left ventral occipito-temporal cortex activation in visual word recognition. PLOS Computational Biology. 2022;18(6):e1009995. doi:10.1371/journal.pcbi.1009995.

7. Olson RK. Dyslexia: nature and nurture. Dyslexia. 2002;8(3):143–159. doi:10.1002/dys.228.

8. Grainger J, Dufau S, Montant M, Ziegler JC, Fagot J. Orthographic Processing in Baboons (Papio papio). Science. 2012;336(6078):245–248. doi:10.1126/science.1218152.

9. Scarf D, Boy K, Uber Reinert A, Devine J, Güntürkün O, Colombo M. Orthographic processing in pigeons (Columba livia). Proceedings of the National Academy of Sciences. 2016;113(40):11272–11276.

10. Linke M, Bröker F, Ramscar M, Baayen H. Are baboons learning ”orthographic” representations? Probably not. PLOS ONE. 2017;12(8):e0183876. doi:10.1371/journal.pone.0183876.

11. Hannagan T, Ziegler JC, Dufau S, Fagot J, Grainger J. Deep Learning of Orthographic Representations in Baboons. PLoS ONE. 2014;9(1):e84843. doi:10.1371/journal.pone.0084843.

12. Hannagan T, Agrawal A, Cohen L, Dehaene S. Emergence of a compositional neural code for written words: Recycling of a convolutional neural network for reading. Proceedings of the National Academy of Sciences. 2021;118(46). doi:10.1073/pnas.2104779118.

13. van Vliet M, Rinkinen O, Shimizu T, Niskanen AM, Devereux B, Salmelin R. Convolutional networks can model the functional modulation of the MEG responses associated with feed-forward processes during visual word recognition. eLife. 2025;13:RP96217. doi:10.7554/eLife.96217.

14. Castles A, Rastle K, Nation K. Ending the Reading Wars: Reading Acquisition From Novice to Expert. Psychological Science in the Public Interest. 2018;19(1):5–51. doi:10.1177/1529100618772271.

15. Gagl B, Hawelka S, Wimmer H. On Sources of the Word Length Effect in Young Readers. Scientific Studies of Reading. 2015;19(4):289–306. doi:10.1080/10888438.2015.1026969.

16. Ziegler JC, Perry C, Zorzi M. Modelling reading development through phonological decoding and self-teaching: implications for dyslexia. Philosophical Transactions of the Royal Society B: Biological Sciences. 2014;369(1634):20120397. doi:10.1098/rstb.2012.0397.

17. Gagl B, Gregorova K, Golch J, Hawelka S, Sassenhagen J, Tavano A, et al. Eye movements during text reading align with the rate of speech production. Nature Human Behaviour. 2022;6(3):429–442. doi:10.1038/s41562-021-01215-4.

18. Yarkoni T, Balota D, Yap M. Moving beyond Coltheart’s N: A new measure of orthographic similarity. Psychonomic Bulletin and Review. 2008;15(5):971–979. doi:10.3758/pbr.15.5.971.

19. Balota DA, Chumbley JI. Are lexical decisions a good measure of lexical access? The role of word frequency in the neglected decision stage. Journal of Experimental Psychology: Human perception and performance. 1984;10(3):340.

20. Fu W, Gagl B. Specifying Precision in Visual-orthographic Prediction Error Representations for a Better Understanding of Efficient Reading. Journal of Cognitive Neuroscience. 2025; p. 1–15.

21. Gagl B, Sassenhagen J, Haan S, Gregorova K, Richlan F, Fiebach CJ. An orthographic prediction error as the basis for efficient visual word recognition. NeuroImage. 2020;214:116727. doi:10.1016/j.neuroimage.2020.116727.

22. Grainger J, Ziegler J. A Dual-Route Approach to Orthographic Processing. Frontiers in Psychology. 2011;volume 2 - 2011. doi:10.3389/fpsyg.2011.00054.

23. Eisenhauer S, Fiebach CJ, Gagl B. Context-Based Facilitation in Visual Word Recognition: Evidence for Visual and Lexical But Not Pre-Lexical Contributions. eneuro. 2019;6(2):ENEURO.0321–18.2019. doi:10.1523/eneuro.0321-18.2019.

24. Taylor J, Davis MH, Rastle K. Mapping visual symbols onto spoken language along the ventral visual stream. Proceedings of the National Academy of Sciences. 2019;116(36):17723–17728.

25. Taylor J, Davis MH, Rastle K. Comparing and validating methods of reading instruction using behavioural and neural findings in an artificial orthography. Journal of Experimental Psychology: General. 2017;146(6):826.

26. Schmalz X, Mulatti C, Schulte-Körne G, Moll K. Effects of complexity and unpredictability on the learning of an artificial orthography. Cortex. 2022;152:1–20.

27. Patzelt EH, Hartley CA, Gershman SJ. Computational Phenotyping: Using Models to Understand Individual Differences in Personality, Development, and Mental Illness. Personality Neuroscience. 2018;1. doi:10.1017/pen.2018.14.

28. Schwartenbeck P, Friston K. Computational Phenotyping in Psychiatry: A Worked Example. eneuro. 2016;3(4):ENEURO.0049–16.2016. doi:10.1523/eneuro.0049-16.2016.

29. Schurr R, Reznik D, Hillman H, Bhui R, Gershman SJ. Dynamic computational phenotyping of human cognition. Nature Human Behaviour. 2024;doi:10.1038/s41562-024-01814-x.

30. Gagl B, Weyers I, Mueller JL. Speechless Reader Model: A neurocognitive model for human reading reveals cognitive underpinnings of baboon lexical decision behavior. In: Proceedings of the Annual Meeting of the Cognitive Science Society. vol. 43; 2021.

31. Rao RPN, Ballard DH. Predictive coding in the visual cortex: a functional interpretation of some extra-classical receptive-field effects. Nature Neuroscience. 1999;2(1):79–87. doi:10.1038/4580.

32. Spratling MW. A review of predictive coding algorithms. Brain and Cognition. 2017;112:92–97. doi:10.1016/j.bandc.2015.11.003.

33. Clark A. Whatever next? Predictive brains, situated agents, and the future of cognitive science. Behavioral and Brain Sciences. 2013;36(3):181–204. doi:10.1017/s0140525x12000477.

34. Olshausen BA, Field DJ. Sparse coding of sensory inputs. Current opinion in neurobiology. 2004;14(4):481–487.

35. Heilbron M, Richter D, Ekman M, Hagoort P, de Lange FP. Word contexts enhance the neural representation of individual letters in early visual cortex. Nature Communications. 2020;11(1). doi:10.1038/s41467-019-13996-4.

36. Heilbron M, van Haren J, Hagoort P, de Lange FP. Lexical processing strongly affects reading times but not skipping during natural reading. Open Mind. 2023;7:757–783.

37. Blank H, Davis MH. Prediction Errors but Not Sharpened Signals Simulate Multivoxel fMRI Patterns during Speech Perception. PLOS Biology. 2016;14(11):e1002577. doi:10.1371/journal.pbio.1002577.

38. Ryskin R, Nieuwland MS. Prediction during language comprehension: what is next? Trends in Cognitive Sciences. 2023;27(11):1032–1052.

39. Eisenhauer S, Gagl B, Fiebach CJ. Predictive pre-activation of orthographic and lexical-semantic representations facilitates visual word recognition. Psychophysiology. 2021;59(3). doi:10.1111/psyp.13970.

40. Ziegler JC, Perry C, Zorzi M. Learning to Read and Dyslexia: From Theory to Intervention Through Personalized Computational Models. Current Directions in Psychological Science. 2020;29(3):293–300. doi:10.1177/0963721420915873.

41. dos Reis M, Thawornwattana Y, Angelis K, Telford M, Donoghue PJ, Yang Z. Uncertainty in the Timing of Origin of Animals and the Limits of Precision in Molecular Timescales. Current Biology. 2015;25(22):2939–2950. doi:10.1016/j.cub.2015.09.066.

42. Pauli J, Gagl B. Understanding the Neuro-Cognitive Mechanisms of Orthographic Learning in Humans and Baboons: A Comparative Study Using Domain-Specific Mechanistic and Domain-General Connectionist Models. bioRxiv. 2025; p. 2025–05.

43. Weinstein B. Matching-from-sample by rhesus monkeys and by children. Journal of Comparative Psychology. 1941;31(1):195–213. doi:10.1037/h0063449.

44. Spinozzi G, De Lillo C, Truppa V, Castorina G. The relative use of proximity, shape similarity, and orientation as visual perceptual grouping cues in tufted capuchin monkeys (Cebus apella) and humans (Homo sapiens). Journal of Comparative Psychology. 2009;123(1):56–68. doi:10.1037/a0012674.

45. Truppa V, De Simone DA, De Lillo C. Short-term memory effects on visual global/local processing in tufted capuchin monkeys (Sapajus spp.). Journal of Comparative Psychology. 2016;130(2):162–173. doi:10.1037/com0000018.

46. Clark W, Colombo M. Seeing the Forest for the Trees, and the Ground Below My Beak: Global and Local Processing in the Pigeon’s Visual System. Frontiers in Psychology. 2022;13. doi:10.3389/fpsyg.2022.888528.

47. Navon D. Forest before trees: The precedence of global features in visual perception. Cognitive Psychology. 1977;9(3):353–383. doi:10.1016/0010-0285(77)90012-3.

48. Fagot J, Tomonaga M. Global and local processing in humans (Homo sapiens) and chimpanzees (Pan troglodytes): Use of a visual search task with compound stimuli. Journal of Comparative Psychology. 1999;113(1):3–12. doi:10.1037/0735-7036.113.1.3.

49. Fagot J, Tomonaga M. Effects of element separation on perceptual grouping by humans (Homo sapiens) and chimpanzees (Pan troglodytes): perception of Kanizsa illusory figures. Animal Cognition. 2001;4(3–4):171–177. doi:10.1007/s100710100109.

50. Burke D, Everingham P, Rogers T, Hinton M, Hall-Aspland S. Perceptual Grouping in Two Visually Reliant Species: Humans (Homo Sapiens) and Australian Sea Lions (Neophoca Cinerea). Perception. 2001;30(9):1093–1106. doi:10.1068/p3239.

51. Deruelle C, Fagot J. Visual search for global/local stimulus features in humans and baboons. Psychonomic Bulletin and Review. 1998;5(3). doi:10.3758/bf03208825.

52. Kurylo DD, Van Nest J, Knepper B. Characteristics of perceptual grouping in rats. Journal of Comparative Psychology. 1997;111(2):126–134. doi:10.1037/0735-7036.111.2.126.

53. Furlan M, Smith AT. Global Motion Processing in Human Visual Cortical Areas V2 and V3. Journal of Neuroscience. 2016;36(27):7314–7324. doi:10.1523/jneurosci.0025-16.2016.

54. Cavoto KK, Cook RG. Cognitive precedence for local information in hierarchical stimulus processing by pigeons. Journal of Experimental Psychology: Animal Behavior Processes. 2001;27(1):3–16. doi:10.1037/0097-7403.27.1.3.

55. Patton TB, Szafranski G, Shimizu T. Male pigeons react differentially to altered facial features of female pigeons. Behaviour. 2010;147(5/6):757–773.

56. Emmerton J, Renner JC. Local rather than global processing of visual arrays in numerosity discrimination by pigeons (Columba livia). Animal Cognition. 2009;12(3):511–526. doi:10.1007/s10071-009-0212-5.

57. Remy M, Güntürkün O. Retinal afferents to the tectum opticum and the nucleus opticus principalis thalami in the pigeon. Journal of Comparative Neurology. 1991;305(1):57–70. doi:10.1002/cne.903050107.

58. Güntürkün O, Hahmann U. Functional subdivisions of the ascending visual pathways in the pigeon. Behavioural Brain Research. 1999;98(2):193–201. doi:10.1016/s0166-4328(98)00084-9.

59. Clark WJ, Colombo M. The functional architecture, receptive field characteristics, and representation of objects in the visual network of the pigeon brain. Progress in Neurobiology. 2020;195:101781. doi:10.1016/j.pneurobio.2020.101781.

60. Fremouw T, Herbranson WT, Shimp CP. Priming of attention to local or global levels of visual analysis. Journal of Experimental Psychology: Animal Behavior Processes. 1998;24(3):278–290. doi:10.1037/0097-7403.24.3.278.

61. Yamazaki Y, Aust U, Huber L, Hausmann M, Güntürkün O. Lateralized cognition: Asymmetrical and complementary strategies of pigeons during discrimination of the “human concept”. Cognition. 2007;104(2):315–344. doi:10.1016/j.cognition.2006.07.004.

62. Martin-Malivel J. Discrimination of contour-deleted images in baboons (Papio papio) and chimpanzees (Pan troglodytes). Animal Cognition. 2011;14(3):415–426. doi:10.1007/s10071-010-0376-z.

63. Dehaene S, Cohen L, Sigman M, Vinckier F. The neural code for written words: a proposal. Trends in Cognitive Sciences. 2005;9(7):335–341. doi:10.1016/j.tics.2005.05.004.

64. Marinus E, de Jong PF. Variability in the word-reading performance of dyslexic readers: Effects of letter length, phoneme length and digraph presence. Cortex. 2010;46(10):1259–1271. doi:10.1016/j.cortex.2010.06.005.

65. Zoccolotti P, De Luca M, Di Pace E, Gasperini F, Judica A, Spinelli D. Word length effect in early reading and in developmental dyslexia. Brain and Language. 2005;93(3):369–373. doi:10.1016/j.bandl.2004.10.010.

66. Sperlich A, Meixner J, Laubrock J. Development of the perceptual span in reading: A longitudinal study. Journal of Experimental Child Psychology. 2016;146:181–201. doi:10.1016/j.jecp.2016.02.007.

67. Snell J, Meeter M, Grainger J. Evidence for simultaneous syntactic processing of multiple words during reading. PLOS ONE. 2017;12(3):e0173720. doi:10.1371/journal.pone.0173720.

68. Weekes BS. Differential Effects of Number of Letters on Word and Nonword Naming Latency. The Quarterly Journal of Experimental Psychology Section A. 1997;50(2):439–456. doi:10.1080/713755710.

69. Yap MJ, Sibley DE, Balota DA, Ratcliff R, Rueckl J. Responding to nonwords in the lexical decision task: Insights from the English Lexicon Project. Journal of Experimental Psychology: Learning, Memory, and Cognition. 2015;41(3):597–613. doi:10.1037/xlm0000064.

70. Bijeljac-Babic R, Millogo V, Farioli F, Grainger J. A developmental investigation of word length effects in reading using a new on-line word identification paradigm. Reading and Writing. 2004;17(4):411–431. doi:10.1023/b:read.0000032664.20755.af.

71. Hawelka S, Schuster S, Gagl B, Hutzler F. Beyond single syllables: The effect of first syllable frequency and orthographic similarity on eye movements during silent reading. Language and Cognitive Processes. 2013;28(8):1134–1153. doi:10.1080/01690965.2012.696665.

72. Ziegler JC, Ferrand L, Jacobs AM, Rey A, Grainger J. Visual and Phonological Codes in Letter and Word Recognition: Evidence from Incremental Priming. The Quarterly Journal of Experimental Psychology Section A. 2000;53(3):671–692. doi:10.1080/713755906.

73. Conrad M, Carreiras M, Tamm S, Jacobs AM. Syllables and bigrams: Orthographic redundancy and syllabic units affect visual word recognition at different processing levels. Journal of Experimental Psychology: Human Perception and Performance. 2009;35(2):461–479. doi:10.1037/a0013480.

74. Galuschka K, Ise E, Krick K, Schulte-Körne G. Effectiveness of Treatment Approaches for Children and Adolescents with Reading Disabilities: A Meta-Analysis of Randomized Controlled Trials. PLoS ONE. 2014;9(2):e89900. doi:10.1371/journal.pone.0089900.

75. Gagl B, Hawelka S, Richlan F, Schuster S, Hutzler F. Parafoveal preprocessing in reading revisited: Evidence from a novel preview manipulation. Journal of Experimental Psychology: Learning, Memory, and Cognition. 2014;40(2):588–595. doi:10.1037/a0034408.

76. Schotter ER, Angele B, Rayner K. Parafoveal processing in reading. Attention, Perception, and Psychophysics. 2011;74(1):5–35. doi:10.3758/s13414-011-0219-2.

77. Gnetov D, Kuperman V. Reading proficiency predicts spatial eye-movement control in the first and second language. Journal of Experimental Psychology: Learning, Memory, and Cognition. 2024;50(8):1315.

78. Sinclair G, Cooley FG, Stringer C, Saunders E, Emmorey K, Schotter ER. Deaf signers target saccades more accurately to the optimal viewing position when reading. Journal of Experimental Psychology: Human Perception and Performance. 2025;.

79. Gagl B, Hawelka S, Hutzler F. A similar correction mechanism in slow and fluent readers after suboptimal landing positions. Frontiers in Human Neuroscience. 2014;8. doi:10.3389/fnhum.2014.00355.

80. Hawelka S, Wimmer H. Visual target detection is not impaired in dyslexic readers. Vision Research. 2008;48(6):850–852.

81. Mason M. Recognition time for letters and nonletters: effects of serial position, array size, and processing order. Journal of Experimental Psychology: Human Perception and Performance. 1982;8(5):724.

82. Ramamurthy M, White AL, Chou C, Yeatman JD. Spatial attention in encoding letter combinations. Scientific Reports. 2021;11(1):24179.

83. Agrawal A, Hari K, Arun S. A compositional neural code in high-level visual cortex can explain jumbled word reading. Elife. 2020;9:e54846.

84. Kakrada E, Colombo M. Beyond the mark: Signatures of self-recognition in fish. Learning Behavior. 2023;52(1):5–6. doi:10.3758/s13420-023-00586-0.

85. Bates D, Mächler M, Bolker B, Walker S. Fitting Linear Mixed-Effects Models, Using lme4. Journal of Statistical Software. 2015;67(1). doi:10.18637/jss.v067.i01.

86. Bates D, Kliegl R, Vasishth S, Baayen H. Parsimonious mixed models. arXiv preprint arXiv:150604967. 2015;.

87. Von der Malsburg T, Angele B. False positives and other statistical errors in standard analyses of eye movements in reading. Journal of memory and language. 2017;94:119–133.

88. Bonferroni C. Teoria statistica delle classi e calcolo delle probabilita. Pubblicazioni del R istituto superiore di scienze economiche e commericiali di firenze. 1936;8:3–62.

